# HeiDI: A model for Pavlovian learning and performance with reciprocal associations

**DOI:** 10.1101/2019.12.18.881136

**Authors:** Robert C. Honey, Dominic M. Dwyer, Adela F. Iliescu

## Abstract

Associative treatments of how Pavlovian conditioning affects conditioned behavior are rudimentary: A simple ordinal mapping is held to exist between the strength of an association (V) between a conditioned stimulus (CS) and an unconditioned stimulus (US; i.e., V_CS-US_) and conditioned behavior in a given experimental preparation. The inadequacy of this simplification is highlighted by recent studies that have taken multiple measures of conditioned behavior: Different measures of conditioned behavior provide the basis for drawing opposite conclusions about V_CS-US_. Here, we develop a simple model involving reciprocal associations between the CS and US (V_CS-US_ and V_US-CS_) that simulates these qualitative individual differences in conditioned behavior. The new model, HeiDI (How excitation and inhibition Determine Ideo-motion), enables a broad range of phenomena to be accommodated, which are either beyond the scope of extant models or require them to appeal to additional (learning) processes. It also provides an impetus for new lines of inquiry and generates novel predictions.

---

*Heidi*, one of the world’s most popular children’s stories, was originally written by Johanna Spyri as two companion pieces: *Heidi: Her years of wandering and learning*, and *Heidi: How she used what she learned*. They describe how Heidi’s predisposition to wander and learn was later evident in her behavior. The central concern of the model that we develop here is the nature of the associative structures that are acquired during Pavlovian conditioning and how these structures result in their behavioral sequelae. Pavlovian conditioning is probably the best-known phenomenon in the history of the scientific study of psychology. The basic procedure and observations can be recounted by people with little or no other knowledge of the field: dogs given pairings of a ringing bell with food come to salivate when the bell rings. HeiDI is a significant revision of the model of Pavlovian conditioning developed by Rescorla and Wagner (1972; Wagner & Rescorla, 1972), and reflects Pavlov’s vision that the study of conditioning provides associative psychology with a scientific basis (Pavlov, 1941, p. 171). Their model has had a profound and enduring influence on the field of animal learning (e.g., Mackintosh, 1975; McLaren, Kaye, & Mackintosh, 1989; Pearce, 1987; Pearce & Mackintosh, 2010; Wagner, 1981), but also on psychology more broadly (e.g., Kruschke, 1992; Gluck & Bower, 1988; Rumelhart, Hinton, & Williams, 1986), and on neuroscience (e.g., Lee et al., 2018; Schultz, Dayan, & Montague, 1997); with 8649 citations at the time of writing this article (Google Scholar). However, the Rescorla-Wagner model offers only the most rudimentary analysis of the associative structures that are acquired during conditioning and how these map onto changes in behavior. Moreover, the model provides no explanation for recent evidence, where different behavioral indices of learning can be taken to support different conclusions about the strength of an association (e.g., Iliescu, Hall, Wilkinson, Dwyer, & Honey, 2018; Flagel, Akil, & Robinson, 2009; Flagel et al., 2011; Patitucci, Nelson, Dwyer, & Honey, 2016). This fundamental problem, together with others that we shall come to (e.g., Miller, Barnet, & Grahame, 1995; Dickinson, Hall, & Mackintosh, 1976; Lubow, 1989; Rescorla, 2000, 2001ab), provided the impetus for the development of HeiDI. The name of the model, HeiDI, reflects the literary reference and links the authors’ surnames to one of the principal issues that the model seeks to address: How excitation and inhibition determine ideo-motion.

## The Rescorla-Wagner Model

The Rescorla-Wagner model proposes that Pavlovian conditioned behavior reflects the formation of an association between the conditioned stimulus (CS) and unconditioned stimulus (US). The presentation of the CS comes to associatively activate the representation or idea of the US and thereby behavior, which can be thus considered ideo-motive: A seemingly reflexive movement effected in response to an idea, in this case the evoked memory of the US. The model has been fundamental to the development of theoretical treatments of associative learning for almost 50 years, and has influenced neurobiological analyses of learning and memory. We briefly review the model here because it provides the principal source of inspiration for the new model that is developed in the remainder of this paper.

According to the Rescorla-Wagner model, the change in the associative strength (ΔV_CS-US_) of a CS on a given trial is determined by the difference between the maximum associative strength supportable by a US (λ) and the pooled associative strength of all stimuli presented on that trial (ΣV_TOTAL-US_). The global or pooled error term (λ – ΣV_TOTAL-US_) allows the model to accommodate phenomena (blocking; e.g., Kamin, 1969; conditioned inhibition; e.g., Rescorla, 1969; contingency effects; e.g., Rescorla, 1968; overshadowing; e.g., Mackintosh, 1978; relative validity; e.g., Wagner, Logan, Haberlandt, & Price, 1968; superconditioning; e.g., Rescorla, 1971) that were beyond the scope of models with separate error terms for each component of a pattern of stimulation (e.g., Bush & Mosteller, 1951; Hull, 1943). It also provides an elegant integration of excitatory conditioning, where the memory of a CS provokes the memory of the US, and inhibitory learning, where a CS can reduce the likelihood of the US memory from becoming active when it otherwise would.

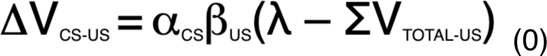

Briefly, the pooled error term means that ΔV_CS-US_ is affected not only by the current associative strength of that stimulus (i.e., V_CS-US_), but also by the presence of other stimuli that have associative strength (i.e., by ΣV_TOTAL-US_). According to the Rescorla-Wagner model, the change in associative strength driven by the discrepancy within the pooled error term (λ – ΣV_TOTAL-US_) is modulated by the product of two learning rate parameters, α_CS_ and β_US_. Rescorla and Wagner (1972) note that “*the value of α roughly represents stimulus salience*” and that “*the assignment of different β values to different USs indicates our assumption that the rate of learning may depend on the particular US employed*”. The two learning rate parameters were confined to the unit interval: 0≤ α_CS_, β_US_ ≤ 1, and enabled the model to capture the fact that the salience of the CS (α_CS_) and nature of the US (β_US_) affect the rate of excitatory learning (see Hall, 1994)^1^. Of particular note, however, is the fact that this model of Pavlovian conditioning did not address – in any systematic fashion – the influence of associative strength (i.e., V) on conditioned responding.

In developing their model and its application to experimental findings, Rescorla and Wagner (1972; p. 77) noted that it was “*sufficient simply to assume that the mapping of Vs into magnitude or probability of conditioned responding preserves their ordering.*”, and that any such mapping would inevitably depend on the details of each experimental situation and on “performance” factors. In a companion paper, when comparing conditioning involving a single CS with conditioning involving a compound of two CSs, they also noted “*that the greater the number of cues which is made available, the more likely it is that the subject will be provided (and perhaps idiosyncratically so) with a single salient cue to which conditioning can rapidly occur*.” (Wagner & Rescorla, 1972; pp. 303-304). This statement acknowledges (parenthetically) the fact that individual differences might affect conditioning (see also, Pavlov, 1941, pp. 373-378), but there has been little appetite to address such differences (empirically or theoretically) and to move beyond simple (group level) assumptions about the translation of learning into performance (see also, for example, Mackintosh, 1975; Miller & Matzel, 1988; Pearce, 1994; Pearce & Hall, 1980; but see, Lesaint, Sigaud, Flagel, Robinson, & Khamassi, 2014; Stout & Miller, 2007). However, there is now evidence demonstrating that the reliance on such assumptions can no longer be sustained; and nor can the idea that Pavlovian conditioning results in unconditioned responses snipped from the US being grafted onto the CS (see Warner, 1932) through a process of stimulus substitution (see Pavlov, 1927; see also, Dwyer, Burgess, & Honey, 2012; Wagner & Brandon, 1989).

## Individual differences

The critical evidence comes from studies of autoshaping in rats, where the brief insertion of a lever (the CS) is immediately followed by the delivery of an appetitive US (e.g., a small quantity of sucrose or a food pellet) into a recessed food well. However, there is no requirement for rats to interact with the signal or enter the food well when the lever is present, but they do. The procedure is an instance of Pavlovian conditioning (see Mackintosh, 1974) and it produces marked individual differences in behavior: Some rats predominantly interact with the lever, others investigate the location where the reinforcer is about to be delivered, and the remainder show patterns of behavior in between these two extremes (e.g., Iliescu et al., 2018; Flagel et al., 2009, 2011; Patitucci et al., 2016; see also, Fitzpatrick et al., 2013; Matzel et al., 2003). Activity directed towards the lever can be measured through recording movements of the lever generated by a rat interacting with it, and is called sign-tracking (e.g., Hearst & Jenkins, 1974; see also, Davey & Cleland, 1982; Timberlake, Wahl, & King, 1982); whereas activity directed towards the food well can be measured by recording occasions when a rat’s snout enters a recess into which reinforcers are delivered, and is called goal-tracking (e.g., Boakes, 1977; Delamater, 1995; Good & Honey, 1991). Both types of behavior can be measured in an automated fashion in conventional experimental chambers. The use of this preparation has highlighted important features of conditioned behavior.

Figure 1 shows the results from a study in which the insertion of one lever was followed by sucrose and the insertion of another (control lever) was not (Patitucci et al., 2016). A median split was used to separate rats into two groups (called sign-trackers and goal-trackers) on the basis of whether their activity during the final block of training (block 6) was predominantly directed towards the lever or food well. This analysis allows the development of the sign-tracking and goal-tracking phenotypes to be traced across training; however, analysis at the level of individual rats reveals that the bias towards sign-tracking or goal-tracking is relatively continuous in nature. The upper panels show the development of lever activity to the lever paired with sucrose and to the control lever followed by no sucrose in the sign-tracking rats (left panel) and goal-tracking rats (right panel). The lower panels show the levels of food well activity across training. When lever activity is used as the assay of discrimination learning, the sign-tracking group show better learning than the goal-tracking group; but when food well activity is used then the reverse is the case. That is, it is not possible to provide a mapping of Vs on to conditioned behavior that provides a coherent interpretation: Focusing on one measure (e.g., sign-tracking) leads to the conclusion that associative learning had proceeded more readily in one set of rats than the other, while focusing on the second measure (e.g., goal-tracking) leads to the opposite conclusion. Even within a preparation, it is not sufficient to assume that there is an ordinal mapping of Vs into the magnitude or probability of conditioned responding. As it stands, the Rescorla-Wagner model is unable to explain why, for any given rat, one response was stronger than the other, and why in some rats goal-tracking was stronger than sign-tracking whereas in other rats this relationship was reversed. That is, it is unable to provide an analysis for why there are both *quantitative* and *qualitative* individual differences in conditioned responding. In fact, these results pose a problem for any theory of learning that assumes a monotonic relationship between a single construct that represents learning and acquired behavior (e.g., Gallistel & Gibbon, 2000; Stout & Miller, 2007).^2^

**Figure 1.**
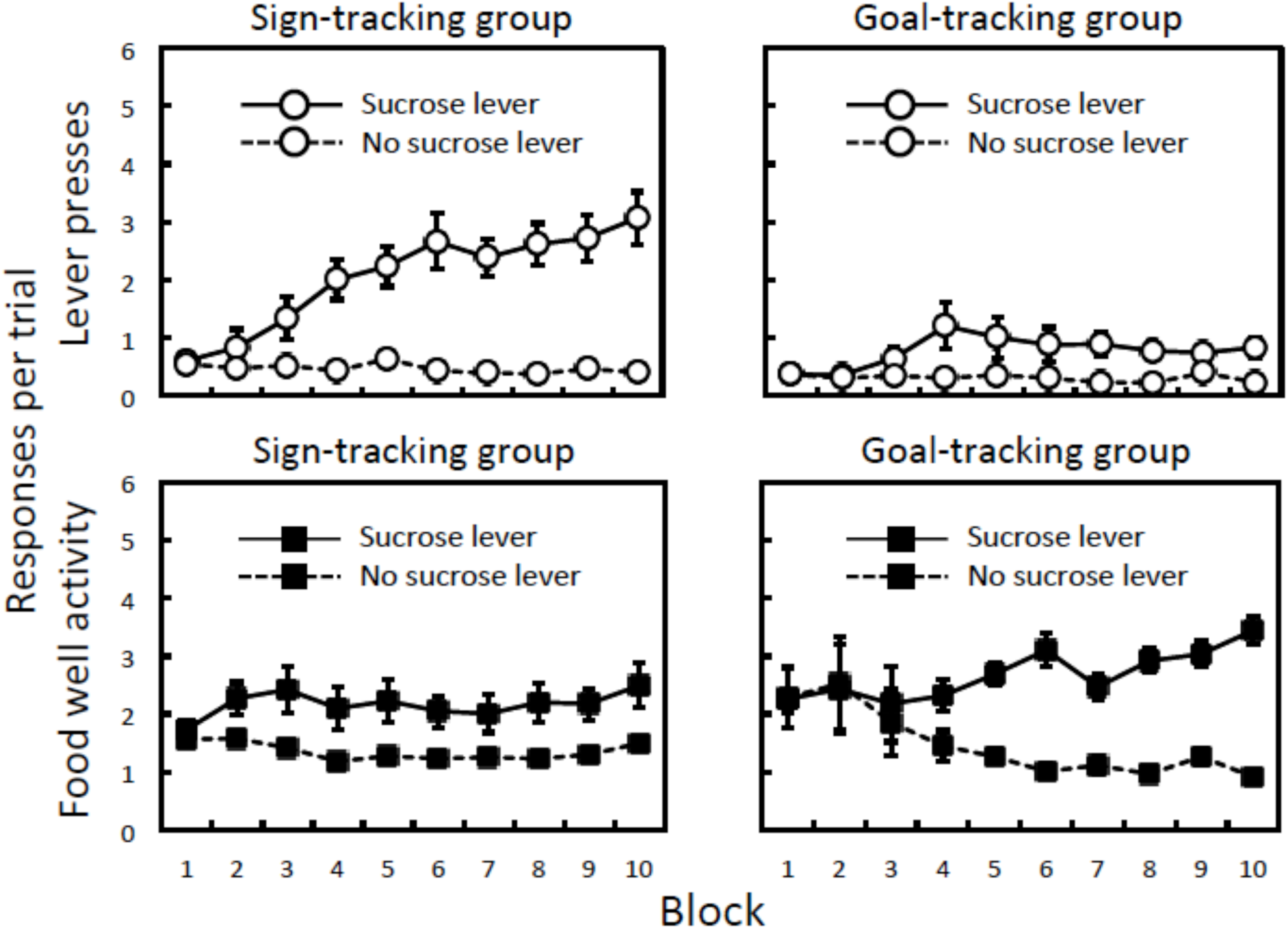
Differences in the form of conditioned behavior. Mean (± SEM) levels of lever activity (sign-tracking) and food well activity (goal-tracking) across 10 training blocks. Rats were divided into sign-trackers (left panels) and goal-trackers (right panels), and the scores are separated for the lever paired with sucrose and the lever that was not. Adapted from: Patitucci, E., Nelson, N., Dwyer, D.M., & Honey, R.C. (2016). The origins of individual differences in how learning is expressed in rats: A general-process perspective. *Journal of Experimental Psychology*: *Animal Learning and Cognition, 42,* 313-324.

## HeiDI: Rationale, architecture and overarching assumptions

The purpose of HeiDI is to offer an account in which the associative structures that are acquired during Pavlovian conditioning are integrated with an analysis of how the knowledge embodied in these structures determines the nature of the responses elicited by a CS, and their relative strengths. In doing so, the model seeks to address challenges to the Rescorla-Wagner model, and other models of Pavlovian learning (e.g., Mackintosh, 1975; Pearce & Hall, 1980; Wagner, 1981).

Figure 2 provides a schematic for the associative structures, to which we will align our analysis of the learning and performance equations that follow. The left-hand panel shows the structure of the model before conditioning has taken place and the right-hand panel shows the structure of the model after conditioning. Before conditioning, the CS is strongly linked to a set of unconditioned responses (r1-r3; e.g., orienting, lever approach, rearing), whereas the US is strongly linked to a set of unconditioned responses (r4-r6; e.g., food well approach, chewing, swallowing). Unconditioned links from the CS to r4-r6 and the US to r1-r3 are assumed to be very weak; and the darkness of the lines between the CS and r1-r6 and between US and r1-r6 denote the relative strengths of these untrained or unconditioned links. In this way, we adopt a general distinction between CS-oriented responses (r1-r3) and US-oriented responses (r4-6; see Holland, 1977, 1984). Importantly, we assume that conditioning results in the formation of reciprocal CS-US and US-CS associations, which are depicted as the presence of dashed lines in the conditioned structure. The general rationale for this assumption, which does not feature in other formal models of Pavlovian conditioning (e.g., Mackintosh, 1975; Pearce & Hall, 1980; Pearce & Mackintosh, 2010; Rescorla & Wagner, 1972), is outlined next. A more specific justification is reserved until the learning rules for these reciprocal associations are presented. We will show that the inclusion of US-CS associations, as well as CS-US associations, provides the basis for HeiDI to explain a wide range of phenomena: In particular, those that have proven difficult to reconcile with the Rescorla-Wagner model (e.g., unequal change in the associative strengths of the components of a compound, Rescorla, 2000; downshift unblocking, Dickinson, Hall, & Mackintosh, 1976) or that have been taken to provide support for models that have emphasized “predictiveness” (e.g., Mackintosh, 1975; Pearce & Hall, 1980; Pearce & Mackintosh, 2010).

**Figure 2.**
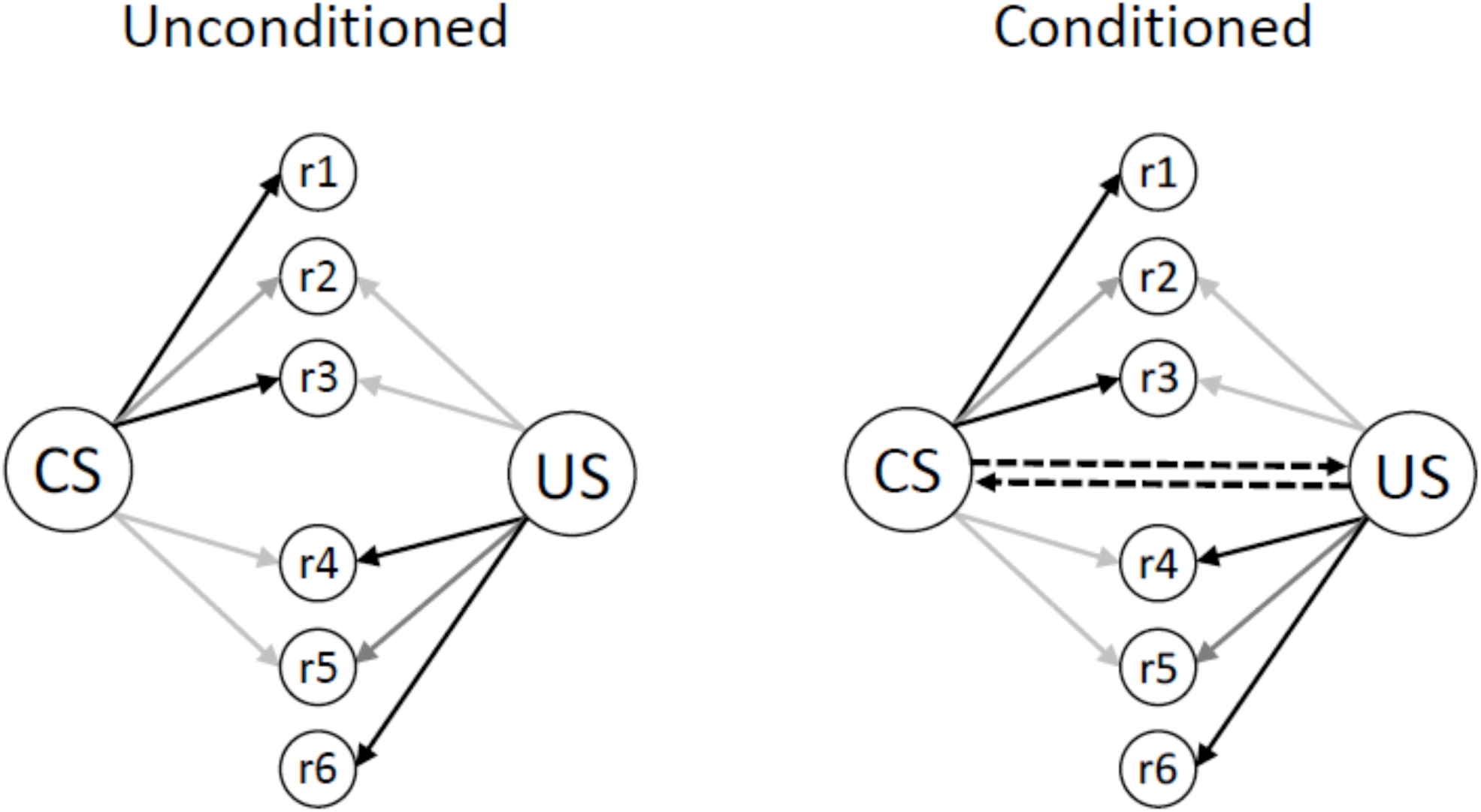
A schematic for the associative structures that underpin the translation of excitatory learning into performance. The left-hand side depicts the model before conditioning (i.e., the unconditioned structure), with the darkness of the arrows indicating the strength of the unconditioned links (i.e., those existing prior to conditioning) between the CS, US and r1-r6. The right-hand side depicts the model after conditioning (i.e., the conditioned structure), which results in changes in the strength of the reciprocal CS-US and US-CS associations between nodes activated by the CS and US (denoted by the dashed lines).

The formation of reciprocal associations between the CS and US creates a functional cell assembly and enables “resonance” between them: When the CS is presented activation propagates to the US, which is propagated back to the CS (e.g., Grossberg, 1980; Hebb, 1949). There is evidence that such reciprocal associations are acquired during forward conditioning in a variety of preparations (e.g., Arcediano, Escobar, & Miller, 2005; Asch & Ebenholtz, 1962; Cohen-Hatton, Haddon, George, & Honey, 2013; Gerolin & Matute, 1999; Honey & Bolhuis, 1997; Honey & Ward-Robinson, 2002; Rescorla & Freberg, 1978; Zentall, Sherburne, & Steirn, 1992); and a complementary literature on the conditions under which US-CS pairings result in conditioned responding to the CS (e.g., Ayres, Haddad, & Albert, 1987; Barnet & Miller, 1996; Cole & Miller, 1999; Heth, 1976; Matzel, Held, & Miller, 1988; Tait & Saladin, 1986). At a theoretical level, in typical Pavlovian conditioning procedures – where the CS precedes but does not co-exist with the US – the memory trace of the CS must be sufficient to support the development of excitatory associations (cf. Wagner, 1981; see also, Barnet & Miller, 1996; Gallistel, 1990; Miller & Barnet, 1993; Silva, Timberlake, & Cevik, 1998). Importantly, while the development of the CS-US association increases the likelihood that the presentation of the CS will activate the US and thereby provoke r4-r6, without the backward associations there would be little change in the likelihood that the CS would provoke r1-r3. The CS-US association allows the presentation of the CS to activate the US node and US-CS association allows activation of the US to increase activation of the CS, which increases the tendency for r1-r3 to become active as a consequence of conditioning.

When a CS is presented, there are two sources of information that are immediately available to an animal upon which performance could be based: The perceived salience of the CS (which is related to α_CS_) and the perceived salience of the US that is activated by the CS (which related to V_CS-US_). A fully effective CS is held to activate the US representation to the value of the perceived salience of the presented US (which relates to β_US_). HeiDI assumes that both of these sources contribute to the nature of performance (cf. Hull, 1949). In particular, the model proposes that the perceived salience of the CS (α_CS_) and the strength of the CS-US association (V_CS-US_) determine how learning is translated into performance through two values, R_CS_ and R_US_. In advance of describing how R_CS_ and R_US_ are calculated exactly, simply assume that increases in α_CS_ results in increases in R_CS_ relative to R_US_ (for a given V_CS-US_ value), while increases in V_CS-US_ results in increase in R_US_ relative to R_US_ (for a given α_CS_ value). Returning to Figure 2, R_CS_ affects behavior via connections from the CS to r1-r6 in Figure 2, and R_US_ affects behavior via connections from the US to r1-r6. We assume that the precise nature of the (alternative) responses generated in a given conditioning preparation will be a function of the interaction between the nature of the CS and US (Holland, 1977, 1984). In the next sections, we first present the learning rules used by HeiDI to determine the development of the reciprocal CS-US and US-CS associations in Figure 1 (Equations 1 and 2); and then provide a simple rule for combining these values upon presentation of the CS (Equation 3). It is worth briefly noting that Equations 1 and 2 reflect the idea that it is the perceived salience of the CS and US, and their associatively generated counterparts, which determine learning. This suggestion is consistent with the idea that individual differences in the perceived salience of the CS and US play a central role in determining individual differences in the expression of learning. We then provide a detailed analysis of how the combined associative strength derived from Equation 3 is separated into two components that affect performance (Equations 4-6). The corresponding simulations of learning and performance are then presented and linked to individual differences in conditioned behavior. Finally, we illustrate how HeiDI provides a natural account for phenomena that challenge the Rescorla-Wagner model, and how it provides alternative analyses for results that have provided the basis for models of Pavlovian learning that include learnt changes in attention or associability (e.g., Mackintosh, 1975; Pearce & Hall, 1980; Pearce & Mackintosh, 2010).

## Learning rules

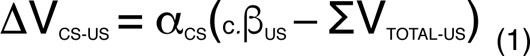

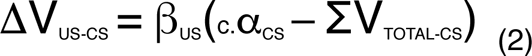

The use of a pooled error term was the central contribution of the Rescorla-Wagner model, allowing it to provide a ready account of the conditions under which excitatory and inhibitory learning occur. HeiDI adopts versions of the pooled error term within Equation 1 and Equation 2, for the formation of CS-US and US-CS associations, respectively. A consolidated list of the HeiDI equations is available at the end of the paper. There is recent evidence that provides direct support for this feature of HeiDI in the context of CS-oriented behavior and US-oriented behavior: A lever CS that provokes sign-tracking can block the acquisition of goal-tracking to an auditory CS, and an auditory stimulus that provokes goal-tracking can block acquisition of sign-tracking to a lever CS (Derman et al., 2018). However, as we shall show, while Equations 1 and 2 incorporate formally equivalent pooled error terms, their functional properties differ when a stimulus compound (AB) is paired with a US: Equation 1 includes a pooled error term that functions as such with respect to the formation of the A-US and B-US associations, whereas in the case of Equation 2 the error term is functionally separate with respect to the formation of the US-A and US-B associations. We will later show how this simple observation enables the use of a pooled error term to be reconciled with results showing that compound (AB) conditioning results in unequal changes in conditioned responding to A and B depending on their prior training histories; an observation that has been considered to implicate separate error terms in Pavlovian conditioning (e.g., Rescorla, 2000; Allman, Ward-Robinson & Honey, 2005; see also, Holmes, Chan, & Westbrook, 2019).

An important feature of Equation 1 is that the perceived salience of the US (relating to β_US_) sets the maximum perceived value of the US retrieved by the CS (relating to V_CS-US_). Similarly, the perceived salience of the CS in Equation 2 (relating to α_CS_) sets the maximum perceived value of the CS retrieved by the US (relating to V_US-CS_). The idea that the perceived salience of directly activated and associatively activated USs influences associative change, receives direct support from results reported by Dwyer, Figueroa, Gassalla, and Lopez (2018). They examined the development of a flavor preference through pairing a flavor CS with an 8% sucrose US. They observed that preceding this concentration of sucrose by either 2% sucrose (generating positive contrast) or 32% sucrose (generating negative contrast) affected the acquisition of the flavor preference: The flavor preference supported by 8% sucrose was larger when it was preceded by 2% sucrose than when it was preceded by 32% sucrose. Moreover, when the changes in the perceived salience of the US (8% sucrose) produced by contrast were directly assessed, through the analysis of licking microstructure, they directly correlated with the size of the resulting preference for the CS flavors.

### Excitatory learning and error correction

Equations 1 and 2 are symmetrical rules governing the formation of CS-US and US-CS associations, respectively. Equation 1 represents a simplification to the Rescorla-Wagner learning rule (Equation 0), and determines the formation of CS-US associations; and Equation 2 provides the formally equivalent rule for US-CS associations. While Equations 1 and 2 include formally equivalent pooled error correcting terms, they have quite different functional properties in conventional conditioning procedures in which a compound of two CSs (AB) precedes a US. In short, the error term in Equation 1 functions as a pooled error term in conventional compound conditioning procedures (Rescorla & Wagner, 1972), whereas the error term in Equation 2 functions as a separate error term in such procedures (Bush & Mosteller, 1951; Hull, 1943). However, it is also worth noting that the model predicts that if a single CS were to be followed by a compound of two USs (US1 and US2), then the association of US1 with the CS would be weaker than if US1 had been paired with the CS in isolation. The prediction that there will be cue competition or overshadowing between the capacities of two USs to become associated with a single CS has received empirical support (e.g., Miller & Matute, 1998).

In Equation 1, α_CS_ is a learning rate parameter confined to the unit interval 0≤ α_CS_ ≤ 1, and c.β_US_ determines the asymptote for the CS-US association; whereas in Equation 2, β_US_ is a learning rate parameter confined to the unit interval 0≤ β_US_ ≤ 1, and c.α_CS_ determines the asymptote for the US-CS association. Note that α_CS_ and β_US_ are dimensionless scalars, but when they serve as the asymptotes for associative strength they are multiplied by a constant of 1 in units of V (c). The requirement for c is that it has units of V in order for the equations to be dimensionally balanced, but the numeric value is not fixed by this requirement. We have assumed c = 1 in units of V for simplicity here. Under these conditions, c.α_CS_ and c.β_US._ will be confined to the unit interval: 0≤ c.α_CS_, c.β_US_ ≤ 1. But, it remains an option for c to take values greater or less than 1 in units of V and in that way for the asymptotic limits of learning to be a multiple of β_US_ in Equation 1 or α_CS_ in Equation 2. When the CS is absent α_CS_ and c.α_CS_ are set to 0 and when the US is absent β_US_ and c.β_US_ are set to 0. In keeping with the Rescorla-Wagner model, α_CS_ and β_US_ are assumed to reflect the perceived salience of the CS and US, respectively. According to Equation 1, the strength of the association from the CS to the US (i.e., V_CS-US_) converges asymptotically on c.β_US_. The change in the strength of the association between CS and the US on a given trial (ΔV_CS-US_) is determined by the error or difference within the pooled error term (c.β_US_ – ΣV_TOTAL-US_); and ΣV_TOTAL-US_ denotes the net associative strength of all of the stimuli presented on that trial. During simple CS-US pairings, excitatory learning ceases when ΣV_TOTAL-US_ = c.β_US_, and the learning rate parameter α_CS_ affects the rate at which V_CS_ approaches c.β_US_. In this case, the pooled error term means that the acquisition of associative strength by a given stimulus will be influenced by the associative strength of other stimuli that accompany it; for example when a compound of two stimuli (A and B) is paired with a US.

Equation 2 is the complementary learning rule governing the formation of the US-CS association. The change in the strength of this association (ΔV_US-CS_) on a given trial is also determined by the discrepancy within the pooled error term (c.α_CS_ – ΣV_TOTAL-CS_); and ΣV_TOTAL-CS_ denotes the associative strength of the US (in typical conditioning procedures). Learning ceases when ΣV_TOTAL-CS_ = c.α_CS_, and the learning rate parameter β_US_ affects the rate at which V_US-CS_ approaches c.α_CS_. Because in typical Pavlovian conditioning procedures there is only one US (cf. Miller & Matute, 1998), the c.α_CS_ value of each CS in a compound (e.g., A and B) sets the asymptote for the association from the US to that CS. This means that the US-CS associations will proceed independently for each of the components of a compound that is paired with a US. That is, while Equation 1 has both the formal and functional properties of Equation (0) and predicts the same phenomena as that model, Equation 2 has equivalent formal properties, but functions in the same way as having a separate error term during compound conditioning (e.g., Bush & Mosteller, 1951; Hull, 1943). How could one test whether the analysis provided by Equations 1 and 2 is accurate?

Consider first the simple case in which two CSs (A and B) are presented together and paired with a US. Under these conditions, the associative strength accrued by A (V_A-US_) and B (V_B-US_) will be less than if these stimuli had been separately paired with the US: An effect known as overshadowing (e.g., Mackintosh, 1978). However, the state of affairs will be different for the reciprocal associations (i.e., V_US-A_ and V_US-B_). They will undergo the same change in associative strength as they would have done had conditioning with each occurred in isolation; because c.α_A_ and c.α_B_ for stimulus A and B set separate asymptotes for the US-A and US-B associations. Of course, the finding that overshadowing is observed under such conditions is uninformative; because Equations 0 and 1 prediction that V_A-US_ will be lower when it has been conditioned in compound with B than when it has been conditioned alone. But, now imagine the same compound conditioning scenario, but that on this occasion a previous stage of training had established A as conditioned excitor (by pairing it with a US) and B had been established as a conditioned inhibitor (by pairing it with the absence of an otherwise predicted US). According to Equations 0 and 1, provided A and B are equally salient (i.e., α_A_ = α_B_) then they should gain equivalent associative strength as a consequence of the AB compound being paired with the US. However, according to Equation 2, while the association between the US and A will not increase (having reached asymptote during the first stage) the association between the US and B will increase, because the US had not previously been paired with B. If the changes in the reciprocal associations were to be combined, then B should have gained greater combined associative strength than A. Rescorla (2000, 2001a; see also, Rescorla, 2001b) has published a series of ingenious experiments that has confirmed this prediction under a variety of circumstances (see also, Allman & Honey, 2005; Allman et al., 2004). We will provide a formal simulation of our analysis of these results, which have been taken to implicate separate error terms in Pavlovian conditioning, once the rules for combining the reciprocal associations have been described, and the way in which associative strength affects performance presented.

As just noted, CS-US pairings create a functional cell assembly through reciprocal associations between the CS and US. To capture this interaction and to simplify our performance rules, it is desirable to combine the net associative strengths of the CS-US association returned by Equation 1 (for V_CS-US_) and the US-CS association returned by Equation 2 (for V_US-CS_). The combined associative strength within this assembly (V_COMB_) is given by Equation 3a.^3^ Here, the reciprocal of the constant, c, is used to translate a value in units of V into a dimensionless value, which means that V_COMB_ has units of V. The combined associative strength of a compound stimulus (V_COMB-AB_) composed of two CSs (A and B) is given by Equation 3b; in which ΣV_AB-US_ is the sum of V_A-US_ and V_B-US_, and V_US-A_ and V_US-B_ are the strengths of the associations between the US and A, and the US and B.

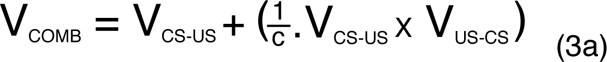

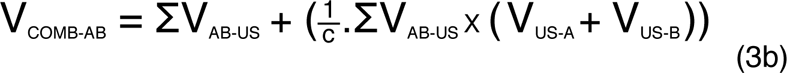

This choice of combination rule recognizes the fact that while the presentation of the CS directly activates the CS-US association, the US-CS association is only indirectly activated by the presentation of the CS. The rule has the general property that the directly activated link in a chain of associations will constrain the impact of the indirectly activated link on performance. In this case, V_CS-US_ will constrain the impact on performance of V_US-CS_. For example, if V_CS-US_ was ≈ 0 and V_US-CS_ was positive, then V_COMB_ ≈ 0 in spite of the fact that the relationship between the CS and US had been encoded (i.e., as V_US-CS_). The significance of this property in the context of HeiDI will become apparent when we consider, in greater detail, the blocking phenomenon (Kamin, 1969).

### Extinction

When conditioning trials with a CS are followed by extinction trials where the CS is presented, but no US occurs, c.β_US_ is set to 0 and ΣV_TOTAL-US_ will be positive. Under these conditions, Equation 1 returns a negative value for ΔV_CS-US_, but Equation 2 returns 0 for ΔV_US-CS_ (because β_US_ = 0). It is worth highlighting this asymmetry between what is learned during conditioning and extinction: excitatory learning involves changes to V_CS-US_ and V_US-CS_, but conventional extinction procedures involve only changes to V_CS-US_. The negative values returned by Equation 1 during extinction can be interpreted in two ways: First, they could denote the growth of negative associative strength (Konorski, 1948; Rescorla & Wagner, 1972; Wagner & Rescorla, 1972). Second, they could denote the formation of an excitatory association between the CS and a ‘No US’ node, which in turn inhibits the US node and thereby reduces in conditioned behavior (see Konorski, 1967; Pearce & Hall, 1980; cf. Zimmer-Hart & Rescorla, 1974). In the first case, the negative values are directly reflected in the underpinning associative structure, and in the second case they reflect the product of an excitatory CS-No US association multiplied by an inhibitory No US–US association. However, according to both interpretations, the net associative strength of the forward association involving the CS (V_CS-US_) is the sum of the positive and negative associative values returned by Equation 1; and the net associative strength of V_US-CS_ is the sum of the positive and negative values returned by Equation 2. Negative values of V_US-CS_ will be returned by Equation 2 when ΣV_TOTAL-CS_ > c.α_CS_. This situation would arise if the US was presented alone after conditioning has taken place or if additional USs were presented in the inter-trial intervals between CS-US pairings. While there is evidence that that is consistent with the prediction that presentations of a US alone after conditioning has taken place can result in a reduction in responding to the CS (see Rescorla, 1973), there is a clear need to test the accuracy of this important prediction from HeiDI across a range of standard conditioning procedures.^4^ In contrast, there is consistent evidence across that adding USs during the inter-trial results in a reduction in conditioned responding to the CS (e.g., Rescorla, 1966, 1968; see also, Durlach, 1983; Gamzu & Williams, 1971, 1973; see also Papini & Bitterman, 1990). In fact, according to HeiDI while both of these manipulations will result in extinction of the US-CS association, adding US presentations during the intervals between CS-US trials will also allow the formation of a context-US association, which should block the development of the CS-US association. In keeping with this analysis, it has been argued that the effects of manipulating CS-US contingency, by adding US alone presentations during conditioning, might be multiply determined (e.g., Baker & Mackintosh, 1979).

Later simulations will confirm the description of the consequences of extinction presented in the previous paragraph. For the time being, it is important to note that according to Equations 1 and 2, extinction leaves a significant contribution to performance completely unchanged (i.e., the US-CS association, V_US-CS_), rather than simply being obscured by additional inhibitory learning, as is the case with the net CS-US association (V_CS-US_). This feature of HeiDI is consistent with the general observation that post-extinction manipulations can reveal the presence of residual excitation in performance, which has represented an ongoing challenge to the Rescorla-Wagner model (e.g., Bouton, 2004). Manipulations that enable the US to be activated (or that disrupt the CS-No US association) will result in a return in performance to the CS.

### Inhibitory learning

If conditioning trials in which stimulus A is paired with a US are intermixed with trials on which A is presented with stimulus B and the US is not delivered, then nonreinforced AB trials will result in a reduction in the net associative strength of A and B will become a net inhibitor. The net associative strength of AB is given by adding the positive and negative values returned by Equation 1 for stimulus A and B. The net associative strength of the US, V_US-CS_, is the sum of the positive and negative associative values returned by Equation 2. According to Equation 2, on nonreinforced AB trials there will be no change in the US-A or US-B associations; again because β_US_ = 0. However, inhibitory learning can also be produced if AB is paired with a US that is smaller in magnitude than the US that is paired with A (e.g., Cotton, Goodall, & Mackintosh, 1982; Nelson, 1987). Under these conditions, β_US_ > 0 and HeiDI predicts that there would be an increase in the *excitatory* strength of the US-A and US-B associations, which would contribute to the values of V_COMB_ for A, B and AB. The prediction that conventional conditioned inhibition training and conditioned inhibition produced by a reduction in reinforcer magnitude should result in different association structures has not been evaluated.

## Performance rules

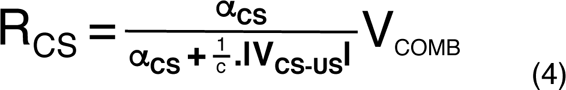

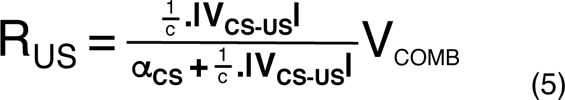

When a CS is presented there are two sources of information that are available to an animal, the perceived salience of the CS (related to α_CS_) and the perceived salience of the US retrieved by through association of the CS with the US (i.e., V_CS-US_), which can be considered an estimate of β_US_ given its relationship with c.β_US_. These two sources of information are held to determine the nature of conditioned behavior. Equations 4 and 5 separate V_COMB_ (derived from Equation 3) into two components: R_CS_ and R_US_. This separation is based on the perceived salience of the CS (i.e., α_CS_) relative to its associative strength (and V_CS-US_). R_CS_ affects behavior via connections from the CS to r1-r6, and R_US_ affects behavior via connections from the US to r1-r6 (see Figure 2). Because in the simulations presented here net V_CS-US_ > 0, the real values of V_CS-US_ can be used to determine R_CS_ and R_US_ in Equations 4 and 5. However, to address the fact that Equation 1 (and Equation 2) can return negative values, the use of absolute values ensures that the proportions in Equations 4 and 5 are ≤ 1. This choice also leaves open the possibility that a net inhibitor could provoke responding when presented alone (cf. Konorski, 1967; Pearce & Hall, 1980), rather than having no effect on performance unless it is presented with an excitor (Konorski, 1948; Wagner & Rescorla, 1972). As before, |V_CS-US_| is transformed into a dimensionless value by multiplying it by 1/c. Because the resulting proportion terms in Equations 4 and 5 are dimensionless, this means that R_CS_ and R_US_ are in units of V. For now, it is sufficient to note that Equation 4 returns a higher value for R_CS_ as the value of α_CS_ increases relative to the value of 1/c.|V_CS-US_|, and Equation 5 returns a higher value for R_US_ as 1/c.|V_CS-US_| increases relative to α_CS_. These two equations are readily extended to accommodate stimulus compounds (AB). To do so, the α values for A and B are simply combined (e.g., added) to form α_AB_, and the net Vs of A and B are combined (e.g., added) to form 1/c.|V_AB-US_|. Similarly, a given stimulus (CS or US) can be conceived of as a set of elements with their own α values and net Vs, which could be entered into Equations 4 and 5 using the same approach (cf. Atkinson & Estes, 1963; see also, Delamater, 2012; Wagner & Brandon, 1989).

While Equations 4 and 5 provide a simple basis for the distribution of the associative properties of the CS-US ensemble (i.e., V_COMB_) to the response-generating units (r1-r6) though R_CS_ and R_US_, they do not specify how these response units once activated affect behavior. One simple possibility is that a given value of R_CS_, for example, results in the same amount of CS-oriented responding (r1-r3) irrespective of the value of R_US_. This possibility equates to there being parallel activation of the response-generating units (r1-r6), and is formally expressed in Equation 6, where R_CS_ and R_US_ are translated into dimensionless values by being multiplied by the reciprocal of the constant, c. According to Equation 6, the activation of a given response unit (e.g., r1) is simply determined by adding the products of (i) multiplying the translated R_CS_ value by the unconditioned link between the CS and r1 (V_CS-r1_), and (ii) multiplying the translated R_US_ value by the strength of connection between the US and the same response unit (e.g., V_US-r1_). We can then make the conventional assumption that the products of Equation 6 (e.g., r1, which is in units of V) are reflected in the overt response (i.e., r1_overt_). That is, the strength of the V_CS-rx_ and V_US-rx_ links scale the associative strengths into observable behavior. There are more complex ways in which R_CS_ and R_US_ might affect r1-r6, involving the interaction between the products of Equation 6 across the set of response-generating units (e.g., McClelland & Rumelhart, 1981). For now, Equation 6 serves as simple placeholder for future theoretical elaboration. However, this level of abstraction does enable future generalization to a range of conditioning preparations. What is more, once the responses and their measurement have been specified it affords quantitative analysis.

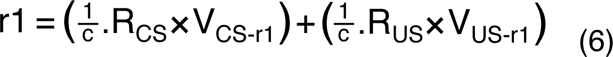

The simulations presented in later sections are derived from Equations 1-5. Equation 6 simply involves multiplying the resulting (dimensionless) R_CS_ and R_US_ values by the fixed strength links between the CS and US nodes and the response units (e.g., the V_CS-r1_ and V_US-r1_ links); with the resulting values being reflect in r_overt_ and their nature being determined by the specific conditioning preparation and responses under consideration. If the two sets of links are equivalent (see Figure 2), then differences in activation of the response units will depend solely on the translated values of R_CS_ and R_US_.

### Individual differences in β_US_

We assume that α_CS_ and β_US_ are fixed for a given CS and US in a given animal, but propose that the perceived salience of the CS (relating to α_CS_) and US (relating to β_US_), and hence α_CS_ and V_CS-US_ in Equations 4 and 5, can vary between animals. This assumption provides the basis for individual differences in R_CS_ and R_US_, because α_CS_ and 1/c.|V_CS-US_| affect the distribution of between CS- and US oriented behavior according to Equations 4 and 5 (remember V_CS-US_ converges on c.β_US_ at asymptote).^5^ This analysis receives support from the observation that rodents who showed a strong liking for sucrose (as measured by licking microstructure; see Dwyer, 2012) are more likely to be goal-trackers (when sucrose was the US) than those who exhibited a weaker liking for sucrose (Patitucci et al., 2016; see also, Morrison et al., 2015). Individual variation in the palatability of sucrose can be aligned to differences in β_US_ that will affect both learning (i.e., the asymptotic value of V_CS-US_ and the rate at which V_US-CS_ reaches asymptote, through Equations 1 and 2) and the distribution of V_COMB_ in performance (through V_CS-US_ in Equations 3-6). As already mentioned, Dwyer et al. (2018) showed that individual differences in the palatability of sucrose (during their experiments involving contrast effects) were positively correlated with the flavor preference learning.

There is additional evidence that is consistent with the proposition that β_US_ for different USs varies between animals, and indeed within a given animal: When separate presentations of two levers are paired with the same US (e.g., food or sucrose) then the bias towards sign-tracking or goal-tracking on one lever correlates with the bias on the other (Iliescu et al., 2018). However, when the presentation of one lever is paired with sucrose and the other lever is paired with food there is no correlation between the biases on the two levers (Patitucci et al., 2016). This pattern of results is consistent with the view that the β_US_ values for two USs (i.e., food and sucrose) can vary between animals and within a given animal (cf. Rescorla & Wagner, 1972).

### Further evidence

A central proposition of HeiDI is that variation in V_CS-US_ (or more precisely 1/c.|V_CS-US_|) interacts with α_CS_ to determine performance. This proposition receives support from the effects of an extinction procedure in which a CS is first paired with a US is then presented alone across a series of trials. Extinction trials should affect net V_CS-US_, conditional on the reduction of c.β_US_ from a positive value to 0 in Equation 1, but no change in α_CS_. The clear prediction is that while both R_CS_ and R_US_ should decrease during extinction (V_COMB_ will reflect the reduction in V_CS-US_; see Equation 3), Equations 4 and 5 predict that this decrease will be less marked for R_CS_ than for R_US_: α_CS_ will remain the same and 1/c.|V_CS-US_| will be lower. Thus, the reduction in V_COMB_ will be partially offset by a rebalancing towards R_CS_ and away from R_US_. This prediction was confirmed in rats that were designated as either sign-trackers or goal-trackers (Ilescu et al., 2018; see also, Ahrens, Singer, Fitzpatrick, Morrow, & Robinson, 2016): In both groups, the tendency for rats to interact with the lever (i.e., sign-tracking) declined less rapidly across extinction trials than did the tendency to interact with the food well (i.e., goal-tracking).

The results from a related conditioning preparation provide converging evidence for the proposed interaction between α_CS_ and 1/c.|V_CS-US_| in determining R_CS_ and R_US_. Kaye and Pearce (1984) gave rats presentations of a localized light that was either paired with the delivery of a food pellet on every trial (in group continuous) or on a randomly scheduled 50% of occasions on which it is presented (in group partial). They observed that when the light was continuously reinforced it maintained a higher level of goal-tracking (food well entries) and a lower level of sign-tracking (orienting and approach to the light) than when the light was partially reinforced (see also, Anselme, Robinson, & Berridge, 2012). According to Equation 1 and 2, net V_CS-US_ will be higher during a continuous than a partial reinforcement schedule, and a continuous reinforcement should result in a greater bias towards goal-tracking (R_US_) and a smaller bias towards sign-tracking (R_CS_) than partial reinforcement, which could result in the opposite bias (see Equations 4 and 5). However, Kaye, and Pearce (1984) also observed that sign-tracking was higher in absolute terms during partial than continuous reinforcement. This finding might reflect the fact that high levels of goal-tracking, during continuous reinforcement, were more likely to interfere (at the level of response output) with sign-tracking than the lower levels of goal-tracking engendered by partial reinforcement (see discussion of Equation 6). In any case, the fact that CS-oriented behavior is maintained by partial reinforcement should also improve an animal’s later ability to detect new relationships involving that CS, a prediction which is supported by evidence from a variety of sources (cf. Pearce & Hall, 1980; Pearce, Wilson, & Kaye, 1988; Swan & Pearce, 1988; Wilson, Boumphrey, & Pearce, 1992; see also, Meyer, Cogan, & Robinson, 2014; Nasser, Chen, Fiscella, & Calu, 2015; Robinson & Flagel, 2009).

## Simulations of learning and performance

### Excitatory conditioning

In all of the simulations that follow, it is assumed that the constant (c) is 1 in units of V. Therefore, the numeric values of α_CS_ and β_US_ are the same as those of c.α_CS_ and c.β_US_, respectively. Figure 3 depicts simulations of the development of the CS-US association derived from Equation 1 (V_CS-US_), the US-CS association derived from Equation 2 (V_US-CS_), and their combined values (V_COMB_) generated by Equation 3. Panels A and B show the simulated values for V_CS-US_, V_US-CS_, and V_COMB_ when α_CS_ was either .30 (panel A) or .70 (panel B) and β_US_ was fixed at .50; and panels C and D show the simulated values for V_CS-US_, V_US-CS_, and V_COMB_ when is α_CS_ was fixed at .50 and β_US_ was either .30 (panel C) or .70 (panel D).

**Figure 3.**
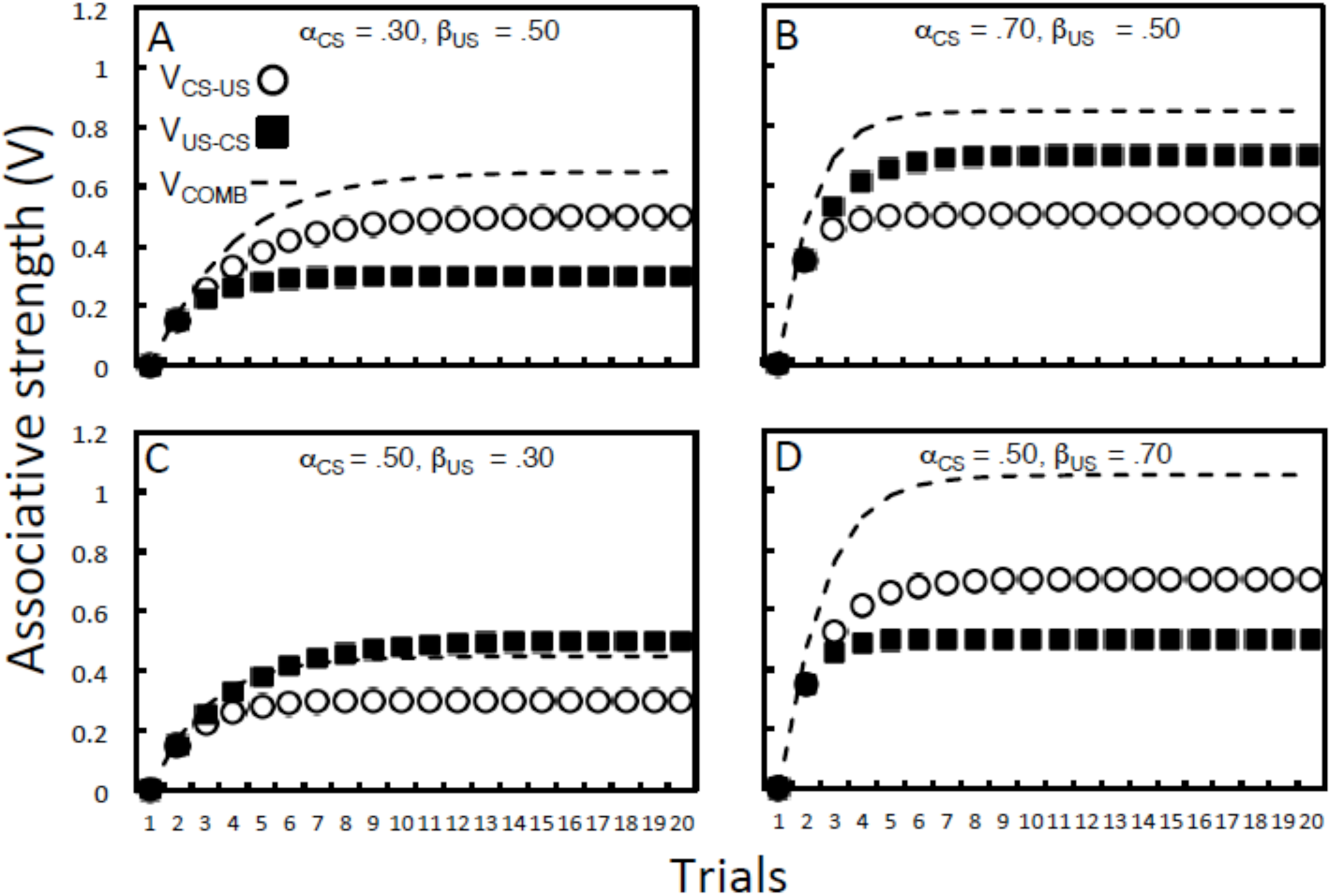
Simulations of associative learning across 20 conditioning trials. Equation 1 was used to generate output values for V_CS-US_, and Equation 2 was used to generate the outputs for V_US-CS_. V_CS-US_ and V_US-CS_ were combined to form V_COMB_ using Equation 3. In panels A and B, α_CS_ was either .30 (A) or .70 (B) and β_US_ was fixed at .50; and in panels C and D, α_CS_ was fixed at .50 and β_US_ was either .30 (C) or .70 (D).

Comparison of panels A and B shows that the CS-US association (V_CS-US;_ open circles) reaches the asymptote derived from the value of β_US_ (i.e., .50) less rapidly when α_CS_ = .30 (panel A) than when α_CS_ = .70 (panel B). Similarly, comparison of the panels C and D confirms that the asymptote for the US-CS association (V_US-CS_; filled squares) derived from the value of α_CS_ (i.e., .50) is reached less rapidly when β_US_ = .30 (panel C) than when β_US_ = .70 (panel D). Finally, the combination of these values (V_COMB_) using Equation 3 is depicted as the hashed line in each panel. Comparison of the adjacent panels (A with C, and B with D) illustrates the impact of the fact that the combination rule (Equation 3) weights V_CS-US_ > V_US-CS_. Thus, in spite of the fact that the summed values of V_CS-US_ and V_US-CS_ are the same in panels A and C (and in B and D), V_COMB_ reflects the fact that V_CS-US_ constrains the impact of V_US-CS_. This fact means that V_COMB_ is higher in panel A than in panel C, and lower in panel B than in panel D.

Figure 4 shows simulations of how Equations 4 and 5 distribute V_COMB_ into R_CS_ and R_US_ values across a series of CS-US pairings. The simulations use the values of V_CS-US_, V_US-CS_ and V_COMB_ taken from Figure 3: Panels A and B use the values of V_CS-US_ and V_COMB_ returned by Equations 1 and 2 when α_CS_ was either .30 (panel A) or .70 (panel B) and β_US_ was fixed at .50; whereas panels C and D use the values of V_CS-US_ and V_COMB_ returned when α_CS_ was fixed at .50 and β_US_ was either .30 (panel C) or .70 (panel D). Maximum V_CS-US_ is determined by c.β_US_ and maximum V_US-CS_ is determined by c.α_CS_. Taking panels A and B first, the asymptote for R_CS_ increased with increases in α_CS_ from .30 (panel A) to .70 (panel B). Comparison of the two panels shows that when β_US_ > α_CS_, R_US_ > R_US_, and when α_CS_ > β_US_, R_CS_ > R_US_. Turning to panels C and D, the asymptote for R_US_ increased with increases in β_US_ from .30 (panel C) to .70 (panel D). Comparison of the two panels shows that when α_CS_ > β_US_, R_CS_ > R_US_, but when β_US_ > α_CS_ then the reverse is the case. The general conclusion is that if α_CS_ = 1/c.|V_CS-US_|, then Equations 4 and 5 distribute V_COMB_ similarly between R_CS_ and R_US_; but if α_CS_ ≠ 1/c.|V_CS-US_| then the distribution of V_COMB_ tracks the component with the largest value (α_CS_ or 1/c.|V_CS-US_|). R_CS_ and R_US_ will affect r1-r6 in the way specified in Equation 6. In general, differences between R_CS_ and R_US_ are simply reflected in the values of V_CS-rx_ and V_US-rx_ for r1-r6, which in turn are reflected in overt responses.

**Figure 4.**
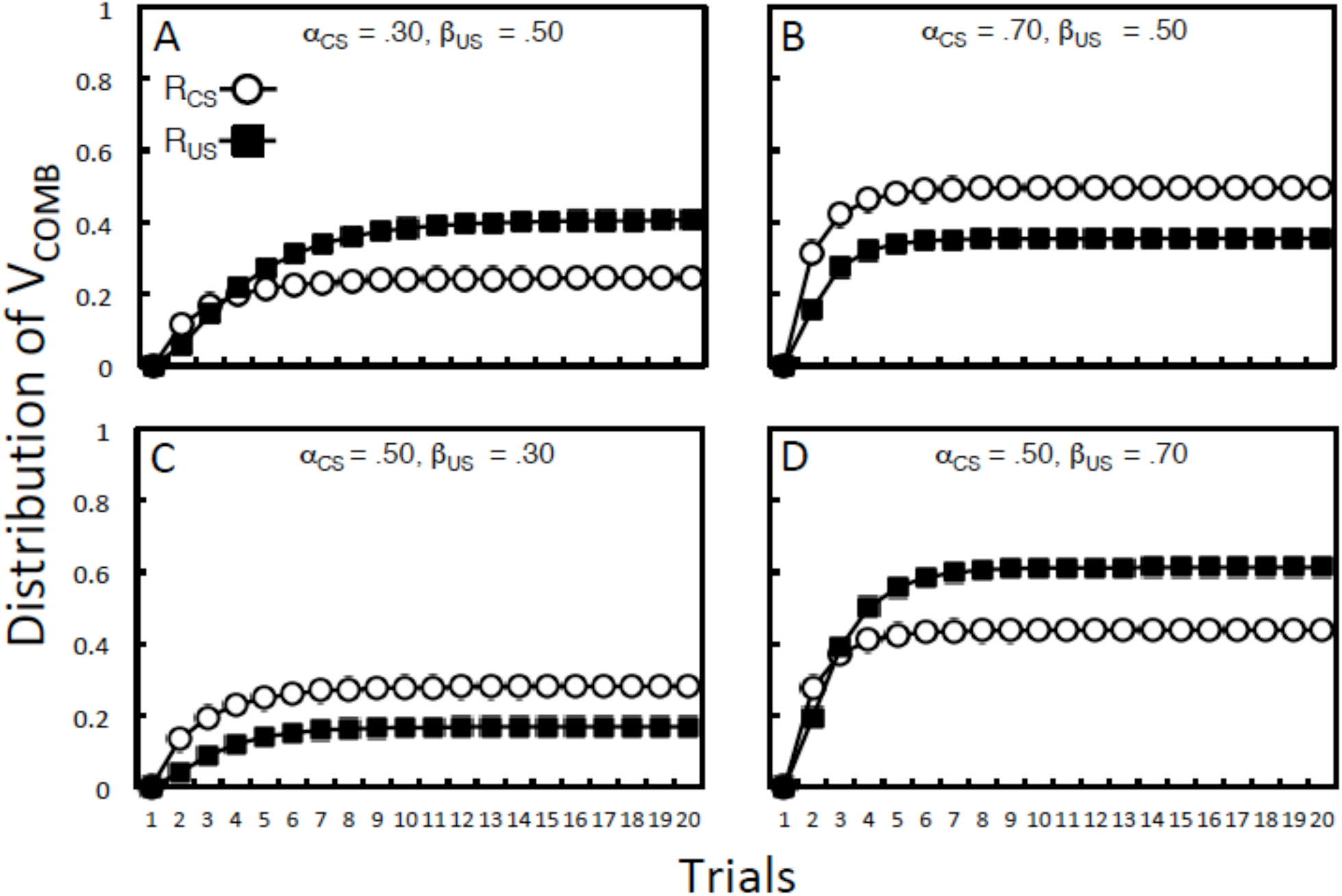
Simulations of the distribution of V_COMB_ into R_CS_ and R_US_ across 20 conditioning trials. R_CS_ and R_US_ outputs were generated using the vales for V_CS-US_, V_US-CS_ and V_COMB_ taken from Figure 3. In panels A and B, α_CS_ was either .30 (A) or .70 (B) and β_US_ was fixed at .50; and in panels C and D, α_CS_ was fixed at .50 and β_US_ was either .30 (C) or .70 (D).

### Extinction

As we have already noted, HeiDI provides a simple analysis of the fact that the CS-oriented component of V_COMB_ (R_CS_) is more persistent during extinction than is the US-oriented component (R_US_; see Iliescu et al., 2018). Briefly, α_CS_ is the same during conditioning and extinction, but net V_CS-US_ declines. Figure 5 shows simulations of conditioning and extinction under conditions in which either R_CS_ > R_US_ during conditioning (panels A and B; α_CS_ = .50 and β_US_ = .30) or R_US_ > R_CS_ (panels C and D; α_CS_ = .30 and β_US_ = .50). Starting with panels A and B, it is clear from panel A that during conditioning V_US-CS_ > V_CS-US_ (when α_CS_ = .50 and β_US_ = .30), and that V_COMB_ is similar to V_US-CS_. During extinction, inspection of panel A shows that V_CS-US_ and V_COMB_ decline, but V_US-CS_ does not (because β_US_ = 0). Panel B shows that the reduction in R_CS_ is (numerically) less marked than R_US_. Moving to panels C and D, during conditioning V_CS-US_ > V_US-CS_ (when β_US_ = .50 and α_CS_ = .30) and V_COMB_ > V_CS-US_. Again, during extinction, V_CS-US_ and V_COMB_ decline, but V_US-CS_ does not. Panel D shows that the reduction in R_CS_ occurs much less rapidly than the reduction in R_US_.

**Figure 5.**
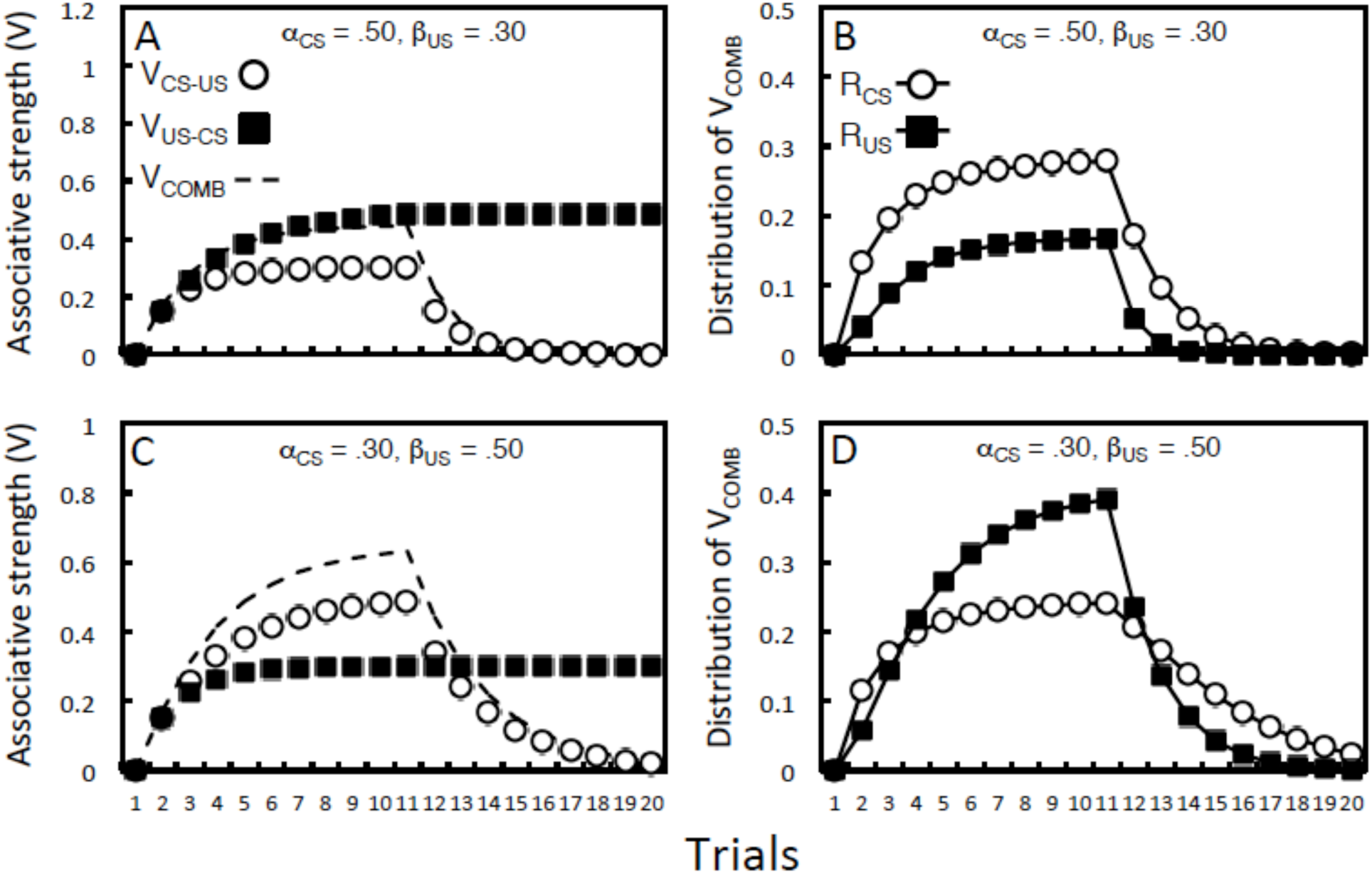
Simulations of conditioning (trials 1-10) and extinction (trials 11-20). Panels A and C depict the output values for V_CS-US_, V_US-CS_ and V_COMB_, and panels B and D show the corresponding output values for R_CS_ and R_US_. The parameters during conditioning were chosen to result in a bias towards R_CS_ (i.e., α_CS_ = .50 and β_US_ = .30; panels A and B) or a bias towards R_US_ (i.e., α_CS_ = .30 and β_US_ = .50; panels C and D). During extinction, β_US_ was set to 0.

### Inhibitory conditioning

Simulations of inhibitory learning, where A is paired with a US and AB is not, were conducted using the same parameters as the simulations for extinction depicted in Figure 5. In this case, both α_A_ and α_B_ were set at .50 and β_US_ = .30, or α_A_ and α_B_ set at .30 and β_US_ = .50. These simulations are shown in Figure 6. V_COMB_ reached a higher asymptote for A, and V_COMB-AB_ for AB took longer to reach asymptote (i.e. ≈ 0), when β_US_ = .50 than when β_US_ = .30 (see panels A and C). When α_A_ and α_B_ were set at .50 and β_US_ = .30 (panel B), the difference between A and AB was more evident in R_CS_ than R_US_, and when α_A_ and α_B_ were set at .30 and β_US_ = .50, the corresponding difference was more evident in R_US_ than R_CS_ (panel D). Other components of this simulation can be explored further using the online app, which returns negative values for inhibitory associative strength using Equations 1 and 2.^6^ This app can also be used to confirm our descriptions of simulations that are not formally presented in the remainder of the paper, and to assess the boundary conditions of our analyses.

**Figure 6.**
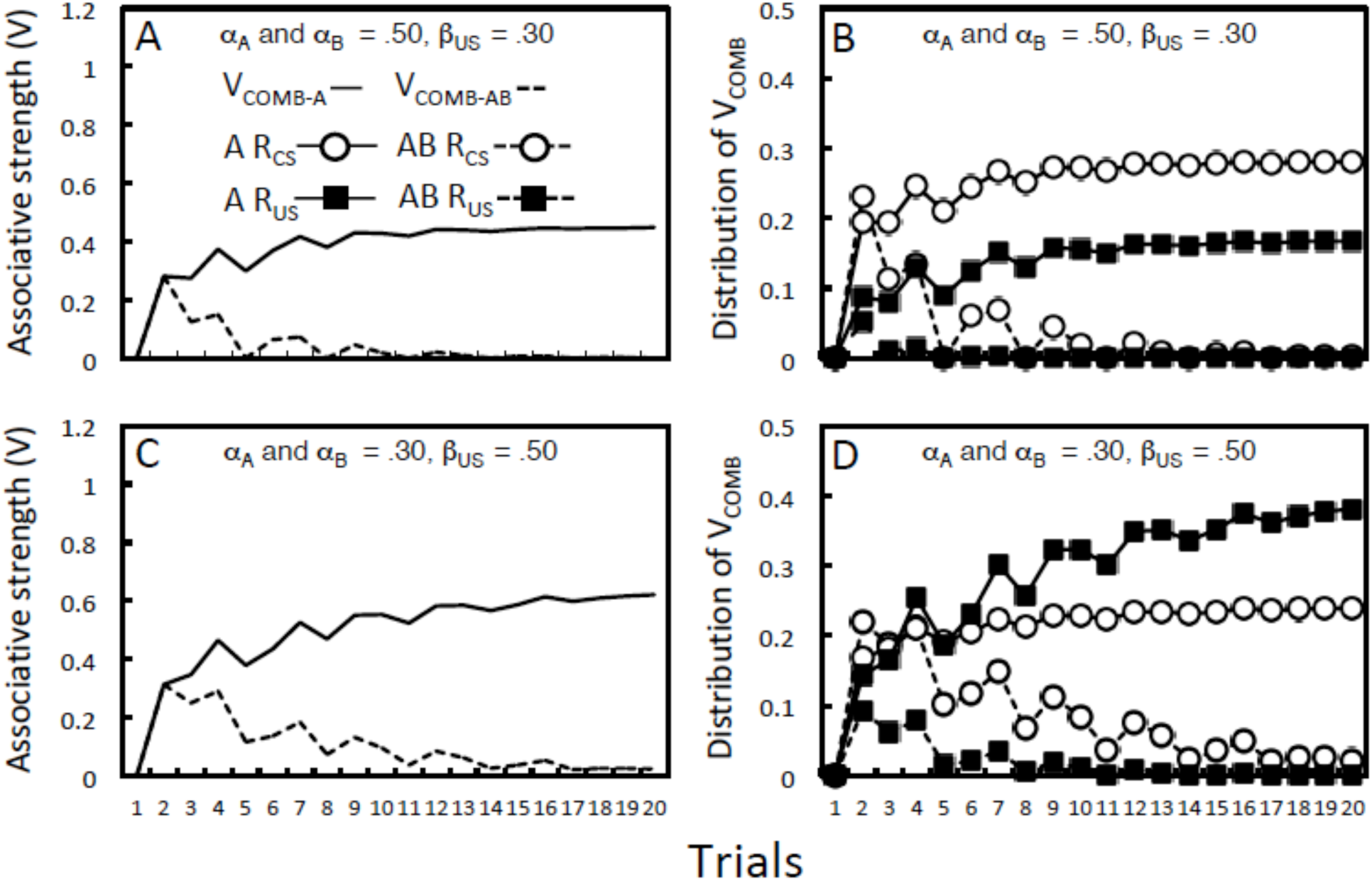
Simulations of conditioned inhibition: A-US and AB-No US. The parameters were chosen to result in a bias towards R_CS_ (i.e., α_CS_ = .50 and β_US_ = .30; panels A and B) or a bias towards R_US_ (i.e., α_CS_ = .30 and β_US_ = .50; panels C and D). Panels A and C depict the output values for V_COMB_ for A and AB, and panels B and D show the corresponding values for R_CS_ and R_US_ for A and AB. The two types of trial were intermixed in a pseudo-random order with the constraint that there were no more than two trials of the same kind in succession. Note that stimulus B acquires net inhibitory properties (not directly shown), but which counteract the excitatory properties that A brings to the AB compound; and that the values for A are taken from the AB trials.

To the best of our knowledge, no experiments have assessed whether the inhibitory properties of stimulus (B) differ depending on whether the excitor with which it was trained (A) evoked CS-oriented (sign-tracking) or US-oriented (goal-tracking) behavior; or indeed whether there are individual differences in how inhibitory learning affects performance. We have used Equations 1-3 in conjunction with Equations 4 and 5 to simulate inhibitory conditioning (e.g., A-food and AB-no food). The simulations confirm that when the α values are higher (i.e., .70) than the net V_CS-US_ supportable by the US (e.g., c.β_US_ = .50), the discrimination between A and AB is more evident (asymptotically) for R_CS_ than R_US_. This effect is evident as R_CS_ being higher than R_US_ for A. They also confirm that when the α values are lower (i.e., .30) than the net V_CS-US_ supportable by the US (e.g., c.β_US_ = .50), the discrimination between A and AB is more evident for R_US_ than R_CS_. This difference is evident as higher R_US_ than R_CS_ for A, and lower R_US_ than R_CS_ for AB. The clear prediction derived from HeiDI is that individual differences in how excitatory learning is exhibited will be correlated with how individual differences in inhibitory learning are manifest. This prediction is novel and its accuracy has yet to be investigated.

We proceed by considering the application on HeiDI to a series of additional phenomena that are central to our understanding of Pavlovian conditioning, but have posed significant challenges to the Rescorla-Wagner model. These phenomena concern: the effects of conditioning a compound with components that have differing associative histories; the effects on performance of combining stimuli with different associative histories; blocking and unblocking; and latent inhibition.

## Compound conditioning and the pooled error term

We have already noted that HeiDI provides a potential reconciliation of the use of a pooled error terms with the observation that stimuli with different associative histories appear to undergo unequal change when they are conditioned in compound. This observation that was taken to be inconsistent with the Rescorla-Wagner model and its successors, which predict equivalent changes provided it is the case that the stimuli are equally salient (see Holmes et al., 2019). To recap: In one set of experiments, Rescorla (2000) first trained two excitors (A and C) and two inhibitors (B and D). Let us assume that A and C both had excitatory associative strength of .50, and B and D both had inhibitory associative strength of −.50 before the compound, AB, was paired with the US (i.e., AB->US). According to Equations 0 and 1, the associative strength of both should increase an equivalent amount: A from .50 to .75 and B from −.50 to −.25. This would mean that the AD compound should have an associative strength of .25 (.75 + −.50) and the BC compound should also have an associative strength of .25 (.50 + −.25). However, according to HeiDI one also needs to consider the fate of the backward associations during compound conditioning: between the US and A, and between the US and B. If we assume that α for all stimuli is .30, then V_US-A_ will be .30 by the end of the first stage of training, but V_US-B_ will be 0, because B has not been paired with the US. This will mean that while V_US-A_ will not change during pairings of AB with the US (the asymptote for V_US-A_ determined by α = .30 will have been reached as a result of the first stage of training), V_US-B_ can increase (e.g., from 0 to .30). Under these conditions, V_COMB-BC_ will higher than V_COMB-AD_. This analysis retains a pooled error term for all associations, but recognizes the fact – hitherto unacknowledged – that associations from the US to A and B will proceed independently of one another in conventional conditioning procedures (i.e., when there is only a single US).

Simulations confirm the accuracy of this analysis across a broad range of parameters, but in the interests of consistency the parameters were set in the way described in the previous paragraph: The α values of A, B, C and D were set at .30; by the end of stage 1, V_A-US_ and V_C-US_ were .50 (i.e., c.β_US_ = .50) while V_B-US_ and V_D-US_ were – .50; and V_US-A_ and V_US-C_ were .30, whereas V_US-B_ and V_US-D_ were 0. Having set these parameters, we then simulated how the CS-US associations involving A and B changed during conditioning with the AB compound (Figure 7A). Inspection of Figure 7A confirms that V_A-US_ and V_B-US_ increased by equivalent amounts, and that while V_US-A_ remained the same, V_US-B_ increased to .30. Figure 7B shows how the associative strengths of AD and BC change when the changes involving A and B were added to the existing strengths of D and B, respectively. Inspection of Figure 7B confirms that the net V_AD-US_ and V_BC-US_ increase equivalently as a consequence of AB conditioning trials (the black symbols overlap with one another). However, while V_US-BC_ increases, V_US-AD_ does not. Figure 7C shows that the V_COMB-BC_ is greater than V_COMB-AD_, reflecting the greater contribution of V_US-BC_ to BC than V_US-AD_ to AD. Finally, Figure 7D reveals that the difference between BC and AD is evident in both R_US_ and R_CS_; but in absolute terms is most evident for R_CS_. This difference reflects the fact that with the parameters employed in the illustrative simulation, the combined alpha scores (α_AD_ and α_BC_ = .60) are greater than the V_AD-US_ and V_BC-US_ (both = .25). When other aspects of the simulation are held constant, but the αs for all stimuli was set at .10 (i.e., α_AD_ and α_BC_ = .20), the absolute difference between BC and AD is (approximately) equally evident for R_US_ and R_CS_.

**Figure 7.**
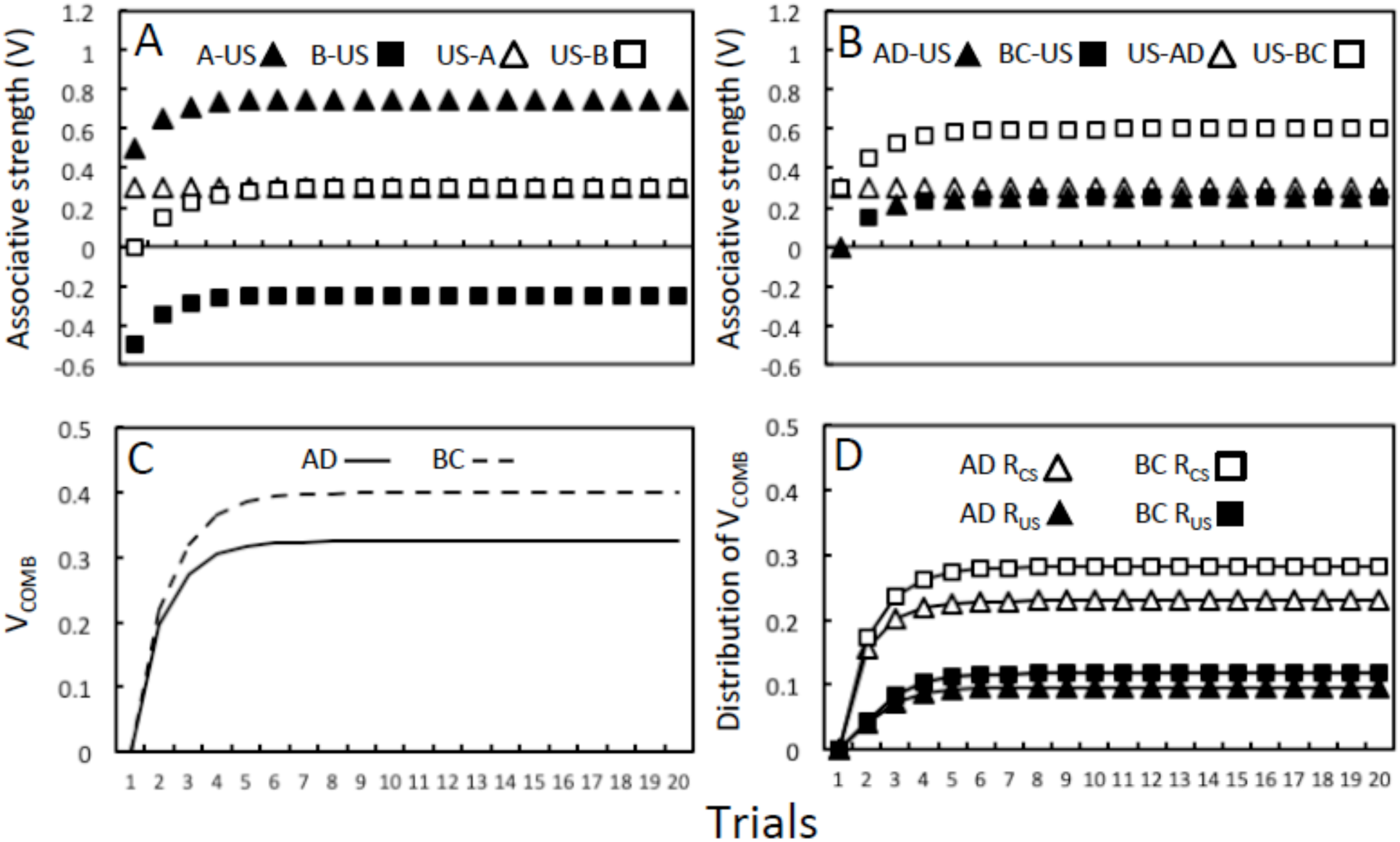
Associative changes when a conditioned excitor (A) and inhibitor (B) are conditioned in compound (AB) and tested with an inhibitor (D) and excitor (C) in compounds AD and BC. Panel A shows the output values for changes in associative strength of the components (A and B) of a stimulus compound (AB) that is paired with a US. Stimulus A (and C) begin compound conditioning with a V_CS-US_ of .50, and V_US-CS_ of .30; whereas B (and D) begin with a V_CS-US_ of −.50 and V_US-CS_ of 0. Panel B depicts the output values for the test compounds: V_AD-US,_ V_US-AD_, V_BC-US_ and V_US-BC_. Panel C shows the output values for the combination of the forward and backward associations for AD (V_COMB-AD_) and BC (V_COMB-BC_), while panel D illustrates how the differences in V_COMB-AD_ and V_COMB-BC_ are reflected in the output values for R_CS_ (CS-oriented behavior) and R_US_ (US-oriented behavior) during the test compounds AD and BC.

## Combining stimuli with different associative histories

Rescorla and Wagner (1972) made the simplifying assumption that the associative strength of a compound stimulus (V_AB-US_) is simply the sum of the individual associative strengths of A and B (i.e., V_A-US_ + V_B-US_). Together with the assumption that V bears an ordinal relationship to performance, the model is constrained to predict that there will be an ordinal relationship between performance to A, B and AB. For example, if two stimuli with excitatory associative strength are combined then performance to the compound AB should exceed both A and B; whereas if one stimulus is excitatory (A) and the other (B) is untrained (and without associative strength) then performance to AB should match A, and both should exceed B. Finally, if A is excitatory and B inhibitory then performance to AB should be less than A and greater than B, unless the excitatory value of A was less than or equal to the inhibitory value of B. While the predictions of HeiDI and the Rescorla-Wagner model mirror one another in some of these cases, they diverge in others.

### Summation

Our analysis begins with the first example, where two CSs (A and B) that have been separately paired with US are predicted to summate when they are combined at test. We used Equations 1 and 2 to generate the requisite individual Vs for stimuli A and B, and Equations 4 and 5 to determine performance. We first confirmed that summation was evident in both R_CS_ and R_US_ irrespective of whether the parameters were chosen to result in a bias towards the R_CS_ (e.g., α_A_ and α_B_ = .50, and β_US_ = .30), or R_US_ (e.g., α_A_ and α_B_ = .50, and β_US_ = .70). However, at an empirical level, summation is not an inevitable consequence of presenting two excitatory stimuli in compound. The circumstances under which summation does and does not occur have yet to be fully determined (Pearce, Aydin, & Redhead, 1997; Pearce, Redhead, & George, 2002), with theoretical analyses tending to focus on how the combination or configuration of stimuli changes the way in which they are processed (e.g., Brandon, Vogel, & Wagner, 2000; Pearce, 1994). For now, we reserve comments about the nature of such ‘configural’ processes for the General Discussion. However, the aforementioned theoretical analyses make an important assumption: Separate conditioning trials with A and B results in them acquiring associative strength (relatively) independently of one another (see Brandon et al., 2000; Pearce, 1994). HeiDI does not make this assumption, and this fact has important implications for the conditions under which summation will be observed.

HeiDI assumes that associations form from the US to the CS. Unlike the development of A-US and B-US associations, which proceed independently of one another, the net US-A association will be weakened on a trial on which B is paired with the US and the net US-B association will be weakened on a trial when A is paired with the US. The extinction of US-A and US-B associations (on B-US and A-US trials respectively) will mean that V_COMB-A_ and V_COMB-B_ would be lower than if A or B were trained alone (i.e., a form of cue interference occurs; cf. Escobar, Matute & Miller, 2001). These facts do not in themselves affect the prediction that summation will be observed (our simulations included these reciprocal associations). However, they do raise the possibility that another form of learning will occur that could constrain summation. To the extent that the A-US and US-B associations enable B to become active on a trial with A, and the B-US and US-A enable A to become active on a trial with B, there is the potential for inhibition to develop between A and B (see McLaren & Mackintosh, 2000; McLaren, Kaye, & Mackintosh, 1989). When A and B are then presented together, the presence of mutual inhibition between them will result in a reduction in their activation, in an analogous fashion to how a conventional conditioned inhibitor affects the ability of a US to become active (cf. Konorski, 1968).

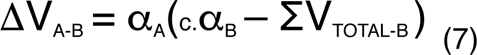

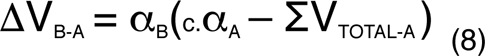

We can first assume that the change in the strength of the association between A and B is governed by Equation 7, and the reciprocal B-A association is governed by Equation 8. These equations are formally equivalent to Equations 1 and 2. They provide a basis for the formation of associations between the elements of a compound (AB), allowing behavior established to one stimulus (e.g., A) to transfer to the other (e.g., B). We will return to these CS-CS associations in the context of a potential analysis of features of blocking and in the General Discussion. The equations also provide the basis for the development of inhibition between A and B when both have been paired with the same US. According to Equations 7 and 8, net inhibition will develop between A and B to the extent that the combined effect of the forward (e.g., A-US) and backward associations (e.g., US-B) provide an indirect basis for V_A-B_ to be positive when B is absent. Thus, on a trial when A is presented, α_B_ = 0 and the ability of A to activate B (i.e., V_A-B_) will depend on multiplying the strengths of the A-US and US-B associations: 1/c.V_A-US_ × V_US-B_; and on a trial when B is presented, α_A_ = 0 and V_B-A_ will depend on: 1/c.V_B-US_ × V_US-A_. The development of this inhibition will mean that when A and B are presented together (e.g., for a summation test) they will be less likely to become active than if they had been presented alone: Performance to an AB compound will be constrained to the extent that inhibition developed between A and B when both are followed by the same US. It is worth noting that such a constraint on summation would be less likely if A and B were to be followed by different reinforcers during conditioning; reinforcers with the same tendency to provoke conditioned responding but with distinct sensory properties (e.g., A-food and B-sucrose).

In keeping with the analysis outlined in the previous paragraph, Watt and Honey (1997) observed that a compound (AB) was more likely to provoke conditioned responding at test if its components had been separately paired with different appetitive reinforcers (food and sucrose) that support the same conditioned response, than if they had been paired with the same reinforcer (food or sucrose; or both food and sucrose, on different trials). In general terms, differences in the development of inhibition between A and B engendered by different training procedures should affect the likelihood of summation being observed. The development of inhibition between A and B, when both are paired with the same outcome, has not been directly assessed in studies of summation or considered at a theoretical level (cf. Brandon et al., 2000; Pearce, 1994). However, there is evidence that is consistent with this suggestion from studies of categorization (Aitken, Bennett, McLaren, & Mackintosh, 1996) and perceptual learning (e.g., Dwyer & Mackintosh, 2002; Mundy, Dwyer, & Honey, 2006).

### External inhibition

When an associatively neutral stimulus (B) is presented with a stimulus with associative strength (A) the conditioned response to that stimulus is often disrupted; an effect known as external inhibition. For example, Pavlov (1927, p. 44) originally observed that the amount of conditioned responding to a CS (in his case the amount of salivation in dogs) was reduced when a stimulus with no associative properties was presented with the CS. This effect is not predicted by the Rescorla-Wagner model, and has been interpreted in terms of a decrease in attention to the CS (Mackintosh, 1974, p. 16). In a set of simulations in which the associative strength of V_B-US_ was set to zero and it was presented with a stimulus (A) that possessed excitatory associative strength (V_A-US_ > 0), the presence of B increased R_CS_ and reduced R_US_ for AB relative to A alone. That is, the predicted effects of adding a neutral stimulus to a CS with excitatory associative strength is to increase the tendency of that associative strength to be evident as CS-oriented rather than US-oriented responding. There is evidence that is consistent with this prediction from studies of a related effect, known as disinhibition. Here, conditioned responding (e.g., instrumental lever pressing for food) can be augmented by the presentation of a stimulus (e.g., a light or white noise; see Brimer & Kamin, 1963; Brimer, 1970). In fact, this effect appears to be most apparent when the level of lever pressing is low (e.g., at the onset of a fixed interval; e.g., Flanagan & Webb, 1962; Hinrichs, 1968; Singh & Wickens, 1969). Unfortunately, none of these studies measured ongoing goal-tracking, which should be the mirror image of behavior directed towards the lever.

### Summation tests for conditioned inhibition

Finally, combining a stimulus with strong excitatory properties (A) and a stimulus with modest net inhibitory properties (B) will mean that V_AB-US_ will take a lower value than V_A-US_. Equations 1 and 2 were used to generate the individual Vs for a reinforced stimulus (A) and a stimulus (B) that was nonreinforced in the presence of A. Equations 3-5 were used to determine the balance between CS- (R_CS_) and US-oriented responses (R_US_). Whether the parameters were chosen to result in a bias towards R_CS_ (e.g., α_A_ and α_B_ = .50, and β_US_ = .30), or R_US_ (e.g., α_A_ and α_B_ = .50, and β_US_ = .70), combining A with B resulted in lower levels of both. The values for R_CS_ and R_US_ for the AB compound would remain positive (albeit lower than those for A alone) because V_COMB_ will still be positive. However, if A had modest excitatory properties and B had strong inhibitory properties, then V_COMB_ would be negative, and as a result R_CS_ and R_US_ would also be negative. Adopting Equation 6 would mean that r1 would be negative (unless either V_CS-r1_ or V_US-r1_ were also negative). In this case, an example of a positive r1 might be to approach the lever and a negative r1 to withdraw from the lever. If the negative values returned by Equations 1 and 2 were construed as involving the activation of a No US node (cf. Konorski, 1967; Pearce & Hall, 1980), then the excitatory V_CS-No US_ association would result in R_CS_ and R_No US_ being positive, and R_No US_ could then directly generate different forms of responding not supported by either the CS or US.

## Blocking: Learning and performance

We noted in the introduction that one of the key features of the Rescorla-Wagner model was its ability to explain how the associative strength of one stimulus within a compound affects the associative strength gained by another stimulus within the compound (e.g., blocking; Kamin, 1969). The formal similarity between Equation 1 and the Rescorla-Wagner model is clear, and like this model, Equation 1 generates these important effects on the development of the CS-US association. However, other features of HeiDI mean that blocking is not – as the Rescorla-Wagner model predicts – inevitable.

In extremis, Equations 1-3 in concert with Equations 4 and 5 provide an account of blocking that is clearly related to the Rescorla-Wagner model: If V_A-US_ ≈ c.β_US_ at the end of a period of training where A has been paired with a US, then conditioning with a compound (AB) will result in little or no increase in the B-US association (i.e., V_B-US_ ≈ 0). However, according to HeiDI, the reciprocal US-B association (V_US-B_) will be unaffected by the fact that A has a reciprocal association with the US (V_US-A_), because the c.α_A_ and c.α_B_ values of A and B provide a separate basis for the formation of these associations. The prediction that the US-B association is not blocked will ordinarily be without consequence because Equation 3 will return a V_COMB_ for B ≈ 0 (i.e., if V_B-US_ ≈ 0 then V_B-US_ + (1/c.V_B-US_ × V_US-B_) ≈ 0). According to Equations 4 and 5, R_CS_ and R_US_ ≈ 0 because V_COMB_ ≈ 0. However, one clear implication of this analysis is that treatments that enable the US-B association to influence performance should reduce the blocking effect; and there is evidence that the performance to a blocked stimulus can be augmented under some conditions (for a review, see Miller et al., 1995; see also, Urcelay, 2017).

Both HeiDI and the Rescorla-Wagner model predict that V_B-US_ (and V_A-US_) will increase during the compound conditioning phase of a blocking procedure if V_A-US_ < c.β_US_. However, unlike the Rescorla-Wagner model, HeiDI predicts that the pattern of performance when B is tested will reflect the values of α_B_ and 1/c.|V_B-US_|. Under these conditions, A might generate US-oriented behavior (when 1/c.|V_A-US_| > α_A_), but the associative strength gained by B might be evident as CS-oriented behavior (when α_B_ > 1/c|V_B-US_|). This simple observation has an important implication: A blocking effect might not be evident if the experimental assay was more sensitive to CS-oriented behavior than to US-oriented behavior. The fact that V_B-US_ is low will reduce V_COMB_ in Equation 3, but its contribution to Equations 4 and 5 (i.e., 1/c.|V_B-US_|) will simultaneously increase the contribution to performance of the CS-oriented component (i.e., R_CS_) and reduce the US-oriented component (i.e., R_US_). While it would be tendentious to argue that failures to observe blocking (e.g., Maes, Boddez, Alfei, Krypotos, D’Hooge, De Houwer, & Beckers, 2016) provide support for the analysis presented above – grounds for such failures abound – there can little doubt that blocking effects can be less complete than a simple rendering of the Rescorla-Wagner model would predict (for a recent review and analysis, see Urcelay, 2017).

However, perhaps the most serious challenge to the account of blocking offered by the Rescorla-Wagner model involves the conditions under which “unblocking” occurs. Conventional procedures for blocking involve two stages in which the reinforcer is the same: A->US and then AB->US. The fact that increasing the number of USs between stage 1 (e.g., A->US1) and stage 2 (AB->US1-US2) results in unblocking (i.e., learning about B) is perfectly consistent with the model, because this change introduces a positive discrepancy in the pooled error term (see Equations 0 and 1). The problematic result is the fact that reducing the reinforcer (i.e., A->US1-US2 and then AB->US1) can also result in responding to B (i.e., unblocking; e.g., Dickinson, Hall, & Mackintosh, 1976). Taken in isolation, Equations 0 and 1 predict that the reduction in the number of reinforcers should have resulted in B acquiring inhibitory properties (e.g., Cotton et al., 1982; Nelson, 1987). ‘Downshift unblocking’, as it is known, has been taken as evidence that the reduction in the US prevents the reduction in attention to B that would ordinarily result from the fact that the US was predicted by A; and allows B to be learnt about (e.g., Mackintosh, 1975; Pearce & Hall, 1980). While there has been some progress in understanding the conditions under which downshift unblocking occurs (Holland, 1988) there is no consensus about its explanation. Many have simply adopted the view that downshift unblocking is *prima facie* evidence that attention can change as a result of experience (Pearce & Mackintosh, 2010). However, a speculative explanation for this effect can be derived from application of HeiDI, without appealing to changes in attention.

The essence of the analysis is that the removal of the second shock allows a within-compound B-A association to form more effectively during downshift unblocking than during standard blocking; and this association allows B to “borrow” the associative properties of A. Consider a blocking procedure in which A is first followed by successive presentations of the same nominal US. We can treat each US as having partially separate representations (US1 and US2). Under these conditions, A will become linked to both US1 and US2 until each link reaches the asymptote determined by c.β_US1_ and c.β_US2_; and critically links will be strengthened between US1 and A, and US2 and A, until their combined associative strength = c.α_A_. When AB is paired with US1 and US2, the associations between B and both US1 and US2 will be blocked; and the combined effect of the US1-A and US2-A associations will mean that B will not be able to enter association with A. However, this will not be the case when US2 is omitted. If we assume that the change in the B-A association is determined by α_B_(c.α_A_ – ΣV_TOTAL-A_), with ΣV_TOTAL-A_ = V_US1-A_ + V_US2-A_ + V_B-A_, then the removal of US2 will enable the strengthening of the B-A association (and further increases in the US1-A association). Under these conditions, downshift unblocking will occur to the extent that the influence of the B-A association in retrieving the associative properties of A with US1 (stronger following downshift unblocking than standard blocking) outweighs the fact that the A-US2 (is weaker) and B-US2 (is negative) after downshift unblocking. This account is speculative, mirroring the fact that our understanding of the conditions under which downshift unblocking occurs remains incomplete (see Holland, 1988). However, it receives support from the results of studies reported by Rescorla and Colwill (1983), which showed that manipulations that should disrupt B-A associations also reduce the difference in performance to B between standard blocking and downshift unblocking.^7^

The simulations presented in Figure 8 for the compound conditioning stage are based – in the interests of simplicity – on the following parameters: α_A_ = α_B_ = .30, and β_US1_ = β_US2_ = .30.^8^ However, the critical difference in the B-A association during standard blocking and downshift unblocking is a general one. At the outset of simulated compound conditioning, for both standard blocking (panels A-C) and downshift unblocking (panels D-F), V_A-US1_ was set to .30 and V_A-US2_ was set to .30 to reflect the assumption that β_US1_ = β_US2_ = .30. Critically, V_US1-A_ and V_US2-A_ were set at .15 for standard blocking, whereas for downshift unblocking V_US1-A_ was set at .15 and V_US2-A_ was set to 0 (to reflect the fact that US2 is absent). For the same reason, V_US2-B_ was also set to 0. Panels A-C (standard blocking) and panels D-F (downshift unblocking) depict the values returned by the combination of Equations 1 and 2 with Equations 7 and 8 for: V_A-US1_, V_US1-A_, V_A-US2_ and V_US2-A_ (panels A and D); V_B-US1_, V_US1-B_, V_B-US2_ and V_US2-B_ (panels B and E); and V_A-B_ and V_B-A_ (panels C and F). Inspection of panels A-C confirms that during standard blocking associations involving A remained the same (panel A), and that associations from US1, US2 and A to B all increased by equivalent amounts (panels B and C). Critically, the B-A association did not develop, and this association can provide no basis upon which B could provoke conditioned responding; and the reciprocal US1-B and US2-B associations cannot – in isolation – contribute to performance. In contrast, during downshift unblocking, because US2 is absent, the US1-A and B-A associations can strengthen (see panels D and E). This will mean both that V_COMB-A_ will be higher following downshift unblocking than standard blocking and that B will be able to access V_COMB-A_ through the B-A association. In order for this state of affairs to generate more performance to B it would need to outweigh the fact that the A-US2 and B-US2 are weaker or inhibitory after downshift unblocking than standard blocking. In the General Discussion, we will consider how associative strength (V_COMB-A_) borrowed by one stimulus (B) from another stimulus (A), with which it has an association (V_B-A_), is manifest in performance. For now, it is sufficient to note that HeiDI provides one formal analysis of how within-compound associations might affect the outcome of blocking and unblocking procedures (cf. Urcelay, 2017).

**Figure 8.**
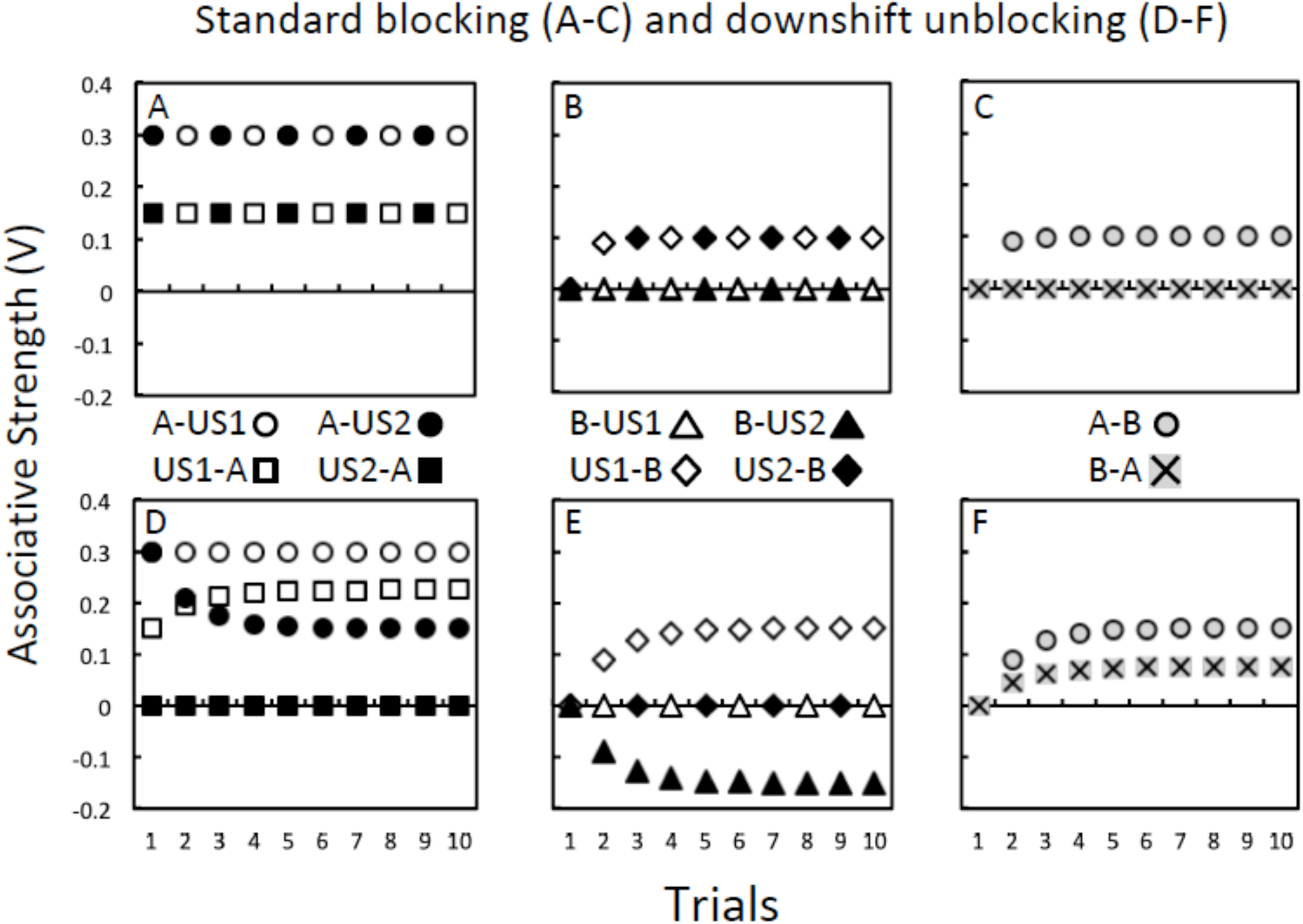
Associative change during compound (AB) conditioning in standard blocking (panels A-C) and downshift unblocking procedures (panels D-E). The parameters used were: α_A_ = α_B_ = .30, and β_US1_ = β_US2_ = .30. At the outset of compound conditioning, A-US1 and A-US2 were set to .30, and US1-A and US2-A were both set to .15. Panels A and D show the output values for the strengths of the A-US1, A-US2, US1-A and US2-A associations returned by Equations 1 and 2 combined with Equations 7 and 8. Note that US2-A is set to 0 in panel D (and US2-B is set to 0 in panel B) to reflect the fact that the US2 is absent; but these associations will not change during unblocking. Panels B and D show the corresponding values for the A-US1, A-US2, US1-A and US2-A associations. Panels C and F show the strength of the A-B and B-A associations. A key observation is that the B-A association gains strength during downshift unblocking (panel F), but not standard blocking (panel C).

## Latent inhibition: An alternative associative analysis

Rescorla and Wagner (1972) recognized the fact that while their model provided a ready account for blocking, it did not address the fact that simple preexposure to a CS retards later excitatory and inhibitory conditioning (for a review, see Hall, 1991; Lubow, 1989). That is, the original model did not provide an account of latent inhibition (Lubow & Moore, 1959). But, why should repeated presentation of a to-be-conditioned stimulus affect the rate at which (excitatory and inhibitory) conditioned performance emerges to that stimulus? This observation in particular, as well as downshift unblocking, has prompted theorists to conclude that models of Pavlovian conditioning need to include another process that changes as a function of experience: attention, associability or CS processing (e.g., Mackintosh, 1975; Pearce & Hall, 1980; Wagner, 1981).

However, a critical feature of latent inhibition, which provides a potential theoretical link with an associative analysis of blocking, is that latent inhibition is context specific. If preexposure to the CS occurs in one context (defined by the cues present in one experimental chamber) and conditioning takes place in another context, then latent inhibition is much reduced (e.g., Grahame, Barnet, Grahame & Miller, 1994; Hall & Honey, 1989; Honey & Good, 1993; Lovibond, Preston, & Mackintosh, 1984; see also, Escobar, Arcediano, Miller, 2002; Wheeler, Stout & Miller, 2004). The general significance of this observation is that it suggests that – during the preexposure stage – animals encode where the stimulus has been presented; for example, by forming a context-CS association (cf. Wagner, 1981). This observation enables HeiDI to provide a simple analysis of latent inhibition: the blocking of the US-CS association by the context-CS association.^9^

We have argued that during excitatory conditioning, performance is determined by both a CS-US association and a US-CS association, and that during inhibitory conditioning, performance could reflect the status of both a CS-No US and a No US-CS association (Konorski, 1968). While a context-CS association will not block the CS-US and CS-No US associations, it will block the development of the US-CS and No US-CS associations. Thus, the simple inclusion of a US-CS association (and No US-CS association) enables an account of latent inhibition that does not require a separate attentional or associability process (e.g., Mackintosh, 1975; Pearce & Hall, 1980) or changes in CS processing of the form envisaged by Wagner (1981).

In addition to this novel analysis of latent inhibition, the presence of a US-CS association means that the effective salience of CSs that are good predictors can be augmented (cf. Mackintosh, 1975). We have demonstrated that the α value of a stimulus affects the rate at which CS-oriented and US-oriented components of performance develop (see Figure 3). The US-CS association provides a natural way in which activation of the US might be reflected back to the CS and maintain its activation. Moreover, we have already noted that when a CS is followed by a reduction in US magnitude (e.g., during extinction or partial reinforcement), CS-oriented responding increases relative to US-oriented responding, which could also affect the subsequent learning involving that CS. HeiDI thereby provides a simple analysis of phenomena that are routinely taken to indicate that the associability of stimuli (their α value) or their processing changes as a result of experience (e.g., Mackintosh, 1975; Pearce & Hall, 1980; Pearce & Mackintosh, 2010; Wagner, 1981). According to our analysis, these phenomena are another product of the reciprocal associations that form between the CS and US, and between the components of stimulus compounds.

## General Discussion

In dispelling out-dated (academic textbook) descriptions of Pavlovian conditioning, Rescorla (1984, p. 151) referred to three primary issues to be addressed in the study of any learning process: “What are the circumstances that produce learning? What is the content of the learning? How does that learning affect the organism’s behavior?”. It is perhaps especially surprising that in the context of Pavlovian learning the final issue – concerning conditioned behavior itself - has become secondary to theorizing directed toward addressing the first two questions. Indeed formal theories of Pavlovian learning have often followed the simplifying stance expressed by Rescorla and Wagner (1972) that it is “*sufficient simply to assume that the mapping of Vs into magnitude or probability of conditioned responding preserves their ordering.*”. The fact that the form of conditioned behavior depends on the nature of both the CS and US (e.g., Holland, 1977, 1984) and that there are marked individual differences in how learning is exhibited (e.g., Iliescu et al., 2018; Patitucci et al., 2016) represent a significant impetus for developing theories that recognize this variety. HeiDI does this.

### Conditions, content and performance

We started by simplifying the Rescorla-Wagner learning rule for forward, CS-US associations, and supplementing it with a formally equivalent rule for reciprocal, US-CS associations (see Equations 1 and 2). The values returned by these equations were then combined (to form V_COMB_) using a rule that weights the associative value of the stimulus that is present (e.g., V_CS-US_) more than an association involving associatively activated nodes (e.g., V_US-CS_; see Equation 3). Finally, when the CS is presented, V_COMB_ is distributed into CS-oriented (R_CS_), and US-oriented (R_US_) components according to the ratio of α_CS_ and 1/c.|V_CS-US_| (see Equations 4 and 5), before being translated into individual responses (see Equation 6). The resulting model, HeiDI, provides the following answers to the three questions posed by Rescorla (1984): (1) On a given trial, learning occurs to the extent that there is a difference between the perceived salience of an event (reflected in β_US_) and the perceived salience of the retrieved representation of that event based on the combined associative strengths of the stimuli presented on that trial (ΣV_TOTAL-US;_ or a difference between c.α_CS_ and ΣV_TOTAL-CS_). (2) Learning is represented in the reciprocal associations between the nodes activated by different stimuli (e.g., CS and US). (3) The combined strength of these reciprocal links (i.e., V_COMB_) is separated into two components (R_CS_ and R_US_) that reflect the perceived salience of the CS (as reflected in α_CS_) relative to the associative strength of the CS (1/c.|V_CS-US_|; which reflects β_US_ through c.β_US_). R_CS_ impacts links between the CS and a set of response units, while R_US_ impacts the links between the US and the same response units. In this way, HeiDI provides a way to capture two classes of conditioned behavior, and individual differences therein, together with the effect of group-level manipulations.

We have highlighted the application of HeiDI to sign-tracking and goal-tracking, which are examples of the general distinction between CS-oriented and US-oriented behaviors. The spatial separation of the two classes of response and the ease with which they are automatically recorded certainly means that they have some methodological advantages over other responses (e.g., those elicited by aversive USs). Nevertheless, we assume that many Pavlovian conditioning procedures result in greater variety in conditioned responses than is routinely measured and used to guide theorizing. We have already illustrated how this practice might complicate interpretation of patterns of results in the case of blocking. However, the two classes of responses that we have considered might themselves be further divided, with the individual elements of the CS and US giving rise to the different responses defined (r1-r6; see Jenkins & Moore, 1973). Expanding HeiDI to accommodate this complexity would not present specific theoretical challenge: with each individual element having its own α or β values and affiliated (unconditioned) responses. However, there are some specific issues that do require further discussion. These involve how associations between the components of a compound stimulus might affect performance, and the nature of the representations of the CS and US.

### Associations between the components of a compound

Conditioned responding to a CS is not only determined by whether it has a direct association with a US. For example, after exposure to a stimulus compound (AB), conditioned responding that is established to B will also be evident when A is presented (e.g., Brogden, 1939; Rescorla & Cunningham, 1978). This effect is known as sensory preconditioning and it is often attributed to the formation of an associative chain that allows A to activate the US through A-B and B-US associations (but see, Lin & Honey, 2016). We have already provided an analysis of how A-B links might form (Equations 7 and 8), and have appealed to such links in providing an analysis of downshift unblocking (cf. Rescorla & Colwill, 1983). The way in which the links in the chain can be combined to determine the level of performance generated by A can be derived from an extension of Equation 3: V_Chain_ = 1/c.V_A-B_ x V_COMB-B_, where V_COMB-B_ = V_B-US_ + (1/c.V_B-US_ x V_US-B_). This formulation means that V_Chain_ < V_COMB-B_ if V_A-B_ < 1. The way in which V_Chain_ is distributed into R_CS_ and R_US_ can be determined using Equations 4 and 5: α_A_ is substituted for α_CS_, 1/c.|V_A-B_ x V_B-US_| is substituted for 1/c.|V_CS-US_|, and V_Chain_ replaces V_COMB_. In terms of the nature of the behavior elicited by A, the most obvious prediction is that it will mirror that evoked by B through direct conditioning (Holland & Ross, 1981). However, according to HeiDI the distribution of CS-oriented and US-oriented behavior will differ between A and B: with CS-oriented responding being more evident (and US-oriented behavior less evident) during A than during B: To the extent that while α_A_ and α_B_ will be the same, 1/c.|V_A-B_ x V_B-US_| < |V_B-US_| (see Dwyer et al., 2012).^10^ This analysis of sensory preconditioning, and of the potential impact of within-compound associations in conditioning procedures more broadly, is relatively straightforward. However, there is another approach to conditioned performance that has also been applied to sensory preconditioning and cue competition effects (e.g., overshadowing and blocking). It deserves consideration because it addresses some of the same issues and phenomena as HeiDI.

The comparator model proposed by Stout and Miller (2007) focuses on how performance to a test stimulus, A, is affected by the stimuli with which it was trained (e.g., B after conditioning with an AB compound). This model builds on the idea that performance to A at test is determined by a comparison between (i) the representation of the US directly retrieved by A, and (ii) the representation of the same US indirectly retrieved by the associative chain: A-B and B-US (see Miller & Matzel, 1988). In this case, B is called the comparator stimulus for A, and following pairings of AB with a US, the tendency for A to generate performance at test is held to be restricted by the fact that its comparator, B, has retrieved a memory of the US. The analysis thereby explains overshadowing and blocking, but also other findings that are problematic for an unreconstructed Rescorla-Wagner model. However, in the case of sensory preconditioning, where AB is first nonreinforced, the model is forced to assume that the fact that B has acquired excitatory associative properties during a second stage increases the potential for A to generate performance. These differing effects of the comparator term (B; termed subtractive and additive) are held to be determined by experience with comparing the US representation retrieved by A with the US representation indirectly retrieved by B. The additive effect occurs when there has been little or no opportunity to experience the two types of retrieved representations (e.g., during simple exposure to AB in sensory preconditioning), and the subtractive effect increases with experience that affords such a comparison (e.g., during multi-trial compound conditioning; pp. 765, Stout & Miller, 2007). In any case, like the Rescorla-Wagner model, the more sophisticated analysis of performance developed by Stout and Miller (2007) provides no ready explanation for the fact that different behavioral measures can provide support for opposing conclusions about how associative strength is translated into performance, which is the focus of interest here. That being said, the fact that within HeiDI the distribution of CS-oriented and US-oriented components of performance reflects the relative values of α_CS_ and 1/c.|V_CS-US_| involves a comparison process of sorts. Certainly, changing the associative strength of stimuli before testing will not only affect V_COMB_, but will also affect R_CS_ and R_US_ through changing 1/c.|V_CS-US_|. As we have already noted, in the context of our previous discussion of blocking, a secure interpretation of the impact of such changes on performance requires behavioral assays that are sensitive to both R_CS_ and R_US_.

### Elemental and configural processes

A final issue, which we mentioned in the section on summation, concerns how models that do not have configural processes address the fact that animals can learn discriminations that are not linearly separable. For example, animals can learn that a tone signals food and a clicker signals no food in one experimental context (a chamber with spotted wallpaper) and the tone signals no food and a clicker signals food in a second context (a chamber with checkerboard wallpaper; e.g., Honey & Watt, 1999). This type of discrimination is interesting because an ‘elemental’ animal – one only capable of representing individual events – should be incapable of learning them: The tone and clicker have the same reinforcement history, as do the spotted and checked chambers, and therefore each of the four combinations or compounds should be equally capable of generating performance. There is an ongoing debate about how different combinations of the same stimuli might be represented in ways that would permit these discriminations to be acquired (e.g., Brandon, Vogel & Wagner, 2000; Pearce, 1994; see also, Honey et al., 2010). For example, different stimulus elements of a given auditory stimulus might become active depending on the context in which they are encountered (e.g., Brandon et al., 2000), or the elements activated by a given pattern of stimulation might come to activate a shared configural representation (e.g., Pearce, 1994; see also, Honey, Close & Lin, 2010). In either case, the elements or configurations thereof (or both, see Honey, Iordanova, & Good, 2014) could be subject to the same learning and performance rules described in Equations 1-6 (see also, Delamater, 2012). However, we should also note that the response units (r1-r6) within the proposed associative architecture for HeiDI (see Figure 2) provide another locus in which combinations of CSs and indeed USs might be represented: The strength of the connections from combinations of CSs and USs to these response units could be modified during conditioning (for a related discussion, see Honey et al., 2010). A formal implementation of the idea that changes in stimulus-response mappings might provide a basis for configural learning is beyond the scope of this article.

### Limitations and further development

We have already noted that Equation 6 provides a simplistic analysis of how changes in R_CS_ and R_US_ might affect performance through their impact on a set of response-generating units (r1-r6). However, taking a step back, what is needed in order to provide a detailed assessment of the accuracy of the predictions that we have derived from HeiDI, is estimates of the perceived salience of both the CS and US on an individual-by-individual basis. Armed with these estimates, we could then provide a quantitative analysis of the fit between predictions of the model and the behavior of animals on an individual basis. We have argued that palatability might provide an estimate of perceived US salience (cf. Patitucci et al., 2016), and one potential estimate of the perceived salience of a CS is the unconditioned orienting behavior that its presentation provokes before conditioning has taken place (cf. Kaye & Pearce, 1984).

### Concluding comments

Pavlovian conditioning has provided a fertile test-bed in which to investigate issues concerning when associative learning occurs, its content, and how it is translated into performance. Of these three issues, formal models have paid least attention to how learning is translated into performance: consideration of performance has been secondary to analyses of the conditions and content of learning. HeiDI begins to redress this imbalance by providing an integrated analysis of all three issues. This analysis could be developed in order to provide a more quantitative analysis, modelling performance at an individual-by-individual level, with the characteristics of the schematic network fully specified. As already noted, it could also be extended to explicitly distinguish between different features of both the CS and the US, which could be tied to different types of response (see Delamater, 2012). In the process of developing this relatively simple model, it has become clear that it is difficult to address one of Rescorla’s three issues without a detailed consideration of the others: developing a more complete understanding of associative learning through the study of Pavlovian conditioning involves multiple constraint satisfaction (Marr, 1982). HeiDI provides general insights into learning, its content and performance that are – at least in part – born out of a more detailed analysis of the variety and individual differences in conditioned behavior. This evidence has been too often neglected, given its theoretical importance and potential translational significance.

## Author note

The underpinning research was conducted when A.F.I. was supported by a School of Psychology PhD studentship, and supervised by R.C.H and D.M.D. All authors contributed to the ideas and preparation of the manuscript for publication. The theoretical ideas contained within this manuscript have not been published in any form; but they formed the basis of a grant funded by the BBSRC UK. The empirical work that underpins the research has been published, but those papers did not contain reference to ideas presented herein. The model was presented at the Associative Learning Symposium at Gregynog (2019), the Australian Learning Group meeting on Magnetic Island (2019), and the Spanish Society for Comparative Psychology in Malaga (2019). These conferences do not have published abstracts or proceedings. We thank John Pearce for his incisive comments on a draft of the paper; and for the reviewers who provided constructive comments that helped to shape the final form of the paper. We also thank Jeremy Hall and Lawrence Wilkinson for their academic input, and for collaborating on the underpinning research through a Strategic Award from the Wellcome Trust (100202/Z/12/Z) on which they were PIs. Correspondence about this article should be addressed to: R.C. Honey;

## Footnotes

1 To enable inhibitory conditioning to occur on trials when the US is absent, Rescorla and Wagner (1972; see also Wagner & Rescorla, 1972) assumed that β takes a positive value when the US is absent but the CS is present; with this value assumed to be lower than on trials when both the CS and US are present. This complexity is avoided in HeiDI.

2 The results presented in Figure 1 have prompted some to argue that sign-tracking and goal-tracking reflect the operation of distinct learning processes. For example, it has been suggested that stimulus-response associations underpin sign-tracking and stimulus-stimulus associations underpin goal-tracking (see Iliescu et al., 2018; Patitucci et al., 2016; see also Lesaint et al., 2014). HeiDI avoids this complexity, and its unmet need to explain when or why distinct learning processes are differentially expressed across animals, because it features a single learning process that is manifest in distinct pathways involving the CS and US.

3 The symmetrical combination rules can be used if the US (rather than the CS) was tested alone (e.g., V_US-CS_ + (1/_C_.V_US-CS_ × V_CS-US_)).

4 There is also evidence that when CS-US pairings are followed by separate presentations of the same US but at a higher intensity (called US inflation) the CR to the CS is amplified (Bouton, 1984; Rescorla, 1974). Under these conditions, in addition to any reduction in net V_US-CS_, presentations of a higher intensity US might change the response units activated by the US, which could affect later performance to the CS.

5 Equations 4 and 5 can be transformed for the case in which the US is presented alone: Under these conditions, β_US_ replaces α_A_ and 1/c.|V_US-CS_| replaces 1/c.|V_CS-US_|.

6 The model and code are implemented as an open source app: https://ynnna.shinyapps.io/HeiDI_model/. The authors can also share Excel spreadsheets that also enable simulations of the critical phenomena to be implemented in a fully transparent manner.

7 It is worth noting that within-compound (A-B) associations could also form during the experiments demonstrating unequal change in the associative strength of the elements of a compound (AB). However, in this case, there was evidence that these associations were not responsible for the effects that were observed (see Allman & Honey, 2005; Rescorla, 2000).

8 The simulations that we report do not include associations between US1 and US2, because they would not influence the formation the excitatory B-A association upon which our analysis rests. Moreover, while the formation of US2-US1 and US1-US2 associations would tend to reduce respectively the A-US1 and A-US2 associations during conditioning with A, the absence of US2 during downshift unblocking would allow increases in both the A-US1 and B-US1 associations. Furthermore, the reductions in the net associative strength of the A-US2 and B-US2 associations produced by the absence of US2 would be less marked than those depicted in Figure 8D and 8E, because US1 would gain a proportion the overall net reduction. In summary, the inclusion of US1-US2 associations increases the likelihood that downshift unblocking would be observed.

9 It should be acknowledged that while the context specificity of latent inhibition is consistent with the view that context-CS associations provide a potential explanation for latent inhibition (and habituation), the fact that attempts to extinguish the context-CS association have often had no effect on latent inhibition is inconsistent with this account (see Baker & Mercier, 1982; Hall & Minor, 1984; but see, Escobar et al. 2002; Grahame et al., 1994; Westbrook, Bond, & Feyer, 1981). However, the interpretation of failures of this kind is not straightforward (see Honey, Good, & Manser, 1998; Honey, Iordanova, & Good, 2010).

10 There is also evidence that the presentation of B itself elicits less responding when it is predicted by A than when it is unpredicted (cf. Wagner, 1981; see Honey, Good, & Manser, 1998; Honey, Hall, & Bonardi, 1993). This observation suggests that there is a refractory period in which, once associatively activated, the presentation of B cannot be fully reactivated.

